# *Arabidopsis* LTR retrotransposons and their regulation by epigenetically activated small RNA

**DOI:** 10.1101/2020.01.24.919167

**Authors:** Seung Cho Lee, Evan Ernst, Benjamin Berube, Filipe Borges, Jean-Sebastien Parent, Paul Ledon, Andrea Schorn, Robert A. Martienssen

**Affiliations:** Howard Hughes Medical Institute, Cold Spring Harbor Laboratory, 1 Bungtown Road, Cold Spring Harbor, NY 11724, USA; Cold Spring Harbor Laboratory, 1 Bungtown Rd, Cold Spring Harbor, NY 11724, USA

**Keywords:** virus-like-particle, retrotransposons, small RNA, transposition

## Abstract

In *Arabidopsis*, LTR-retrotransposons are activated by mutations in the chromatin remodeler DECREASE in DNA METHYLATION 1 (DDM1), giving rise to 21-22nt epigenetically activated siRNAs (easiRNAs) that depend on RNA DEPENDENT RNA POLYMERASE 6 (RDR6). We purified virus-like-particles (VLPs) from *ddm1* and *ddm1rdr6* mutants in which genomic RNA is reverse transcribed into complementary DNA. Next generation short-read and long-read sequencing of VLP DNA (VLP DNA-seq) revealed a comprehensive catalog of active LTR-retrotransposons without the need for mapping transposition, and independent of genomic copy number. Linear replication intermediates of a functionally intact copia element *EVADE* revealed multiple central polypurine tracts (cPPT), a feature shared with HIV where cPPT promote nuclear localization. For one member of the *ATCOPIA52* subfamily (*SISYPHUS*), cPPT intermediates were not observed, but abundant circular DNA indicated transposon “suicide” by auto-integration within the VLP. easiRNA targeted *EVADE* genomic RNA, polysome association of *GYPSY* (*ATHILA*) subgenomic RNA, and transcription via histone H3 lysine-9 dimethylation. VLP DNA-seq provides a comprehensive landscape of LTR-retrotransposons, and their control at transcriptional, post-transcriptional and reverse transcriptional levels.

## Introduction

Long terminal repeat (LTR) retrotransposons and endogenous retroviruses are a major component of the large genomes of most animal and plant species (Huang et al., 2012; Wang et al., 2014). The mouse genome, for example, contains more than one million endogenous retroviruses, of which only a handful are autonomous elements (Gagnier et al., 2019; Huang et al., 2012). In *Arabidopsis thaliana,* ancient Ty3/gypsy type *ATHILA* elements comprise ~3% of the genome, mostly in pericentromeric regions, and relatively young Ty1/copia *ATCOPIA* elements (~1% of the genome) are often found in euchromatic regions (Marco and Marin, 2008; Pereira, 2004). Inhibition of retrotransposons by small RNA has been reported in metazoans and plants, as well as in fission yeast, and occurs at the transcriptional and post-transcriptional levels. In Drosophila, piwi-interacting RNA (piRNA) trigger transcriptional silencing of transposons in the germline (Czech et al., 2018), resembling fission yeast in this respect (Volpe et al., 2002). By contrast, *Ago2* and *Dcr2* lie in the small interfering RNA pathway, and their mutation results in increased somatic retrotransposition (Xie et al., 2013). In mammalian embryos, 3’ tRNA fragments (3’-tRF) control transposition of LTR retrotransposons both post-transcriptionally and by direct inhibition of reverse transcription (Schorn et al., 2017). In *Arabidopsis*, transcriptional activation of Ty1/copia retrotransposons is triggered by stress or loss of DNA methylation, and their retrotransposition rates are enhanced by loss of the 24nt small RNA pathway (Ito et al., 2011; Mirouze et al., 2009). By contrast, in *ddm1* mutants and wild-type pollen, most transposons are transcriptionally active and their RNA transcripts are processed into 21-22nt easiRNA (Lippman et al., 2004; Slotkin et al., 2009). In *ddm1* mutants, easiRNA are generated by RDR6 and diverse miRNA (Creasey et al., 2014; Nuthikattu et al., 2013) from the non-functional *ATHILA2* and *ATHILA6* Ty3/gypsy retrotransposons but also from the functional TY1/copia element *EVADE*. In wild-type, retroelements generate easiRNA only in pollen, where they are targeted at the primer binding site (PBS) by miR845, and biogenesis occurs via a non-canonical pathway (Borges et al., 2018).

Ty1/copia elements in plants have a single open reading frame that encodes both the capsid protein (GAG), and the polyprotein (POL) comprised of reverse transcriptase, RNase H, and integrase. These proteins co-assemble with their genomic RNA (gRNA) into virus-like particles (VLPs), the cytoplasmic compartments encapsulated by GAG in which retrotransposon cDNA intermediates are produced (Supplemental Fig. S1) (Finnegan, 2012; Pachulska-Wieczorek et al., 2016; Peterson-Burch and Voytas, 2002; Sabot and Schulman, 2006). Ty3/gypsy elements also have a single GAG-POL ORF, although the POL proteins are in a different order. In both Drosophila and plants, the Ty1/copia GAG protein is translated from an abundant, alternatively spliced subgenomic RNA (Chang et al., 2013; Yoshioka et al., 1990). In *Arabidopsis*, Ty1/copia elements, the subgenomic *GAG* RNA is more efficiently translated than unspliced GAG-POL transcripts, and blocking splicing leads to significant reduction of GAG protein translation (Oberlin et al., 2017).

After VLP formation in the cytoplasm, LTR retrotransposons proliferate through tRNA-primed reverse transcription of gRNA, followed by nuclear import of double-stranded cDNA and integration into new loci (Chapman et al., 1992; Schorn and Martienssen, 2018) (Supplemental Fig. S1). In yeast and *Arabidopsis*, tRNA-iMet initiates reverse transcription of the LTR from the PBS to the 5’ end of the R region making minus-strand strong-stop DNA (Chapman et al., 1992; Griffiths et al., 2018; Mules et al., 1998; Schorn and Martienssen, 2018). RNase H degrades the template RNA upstream of the PBS, and minus-strand strong-stop DNA is transferred to the 3’ LTR to prime minus strand cDNA synthesis toward the PBS (Supplemental Fig. S1). During the extension of minus-strand cDNA synthesis, the template RNA is degraded except for an RNase H-resistant polypurine tract (PPT) near the 3’ LTR (Wilhelm et al., 2001). This PPT RNA fragment primes plus-strand strong-stop DNA synthesis up to U5 and the PBS sequence from the translocated minus strand (Supplemental Fig. S1B). Then, the plus-strand cDNA is transferred to the 5’ end to prime full-length double-stranded DNA. Additional central PPT (cPPT) can also initiate plus-strand synthesis which is displaced by the 3’ end of plus-strand DNA causing DNA flaps to form during Ty1 replication (Garfinkel et al., 2006). cPPT and DNA flaps have been found in the lentivirus HIV-1 where they play roles in nuclear import and in preventing mutagenesis by the cytidine deaminase APOBEC (Hu et al., 2010; VandenDriessche et al., 2002; Wurtzer et al., 2006; Zennou et al., 2000).

The capacity for specific retrotransposons to proliferate by integration into new genomic loci can only be confirmed by observing these insertions using transposition assays, whole genome sequencing in an active background, or by comparing insertion sites among individuals within a population. However, a number of methods including mobilome-seq (Lanciano et al., 2017) and ALE-seq (Cho et al., 2019) have been developed to catalog elements that may be transpositionally competent by sequencing transposition intermediates prior to integration. Short reads from mobilome-seq cover specifically circular extrachromosomal DNA (ecDNA) which arise from recombination of linear ecDNA. ALE-seq generates reads covering only the 5’-LTRs of linear ecDNA and depends on the use of reverse transcription primers specific to the PBS of the elements being queried. These linear intermediates are more directly predictive of transposition competence, as circular products do not undergo chromosomal integration (Lanciano et al., 2017; Sloan and Wainberg, 2011). We have developed an alternative strategy that enables comprehensive sequencing of products from isolated VLPs. VLPs have been isolated in yeast and Drosophila (Bachmann et al., 2004; Eichinger and Boeke, 1988; Kenna et al., 1998) as well as in plants (Bachmair et al., 2004; Jaaskelainen et al., 1999), but sequencing of their cDNA contents has not been reported. Our method captures linear products as well as abortive linear and circular intermediates missed by other approaches. By sequencing these intermediates from different genetic backgrounds, insights can be gained into mechanisms of genetic and epigenetic regulation.

In this study we examined multiple layers of LTR retrotransposon control in *Arabidopsis*, utilizing *ddm1* mutants where transposons are transcriptionally active and *ddm1rdr6* mutants deficient in easiRNA (Creasey et al., 2014; Lippman et al., 2004; Vongs et al., 1993). We sought to determine whether easiRNA act to silence retroelements at transcriptional and post-transcriptional levels using genome-wide polysomal RNA (translatome), chromatin immunoprecipitation (ChIP), and small RNA sequencing. Furthermore, we performed VLP DNA sequencing to capture the full complement of cDNA intermediates generated during retrotransposition for the first time, and to evaluate whether their sequence features are diagnostic of functional potential.

## Results

### Characterization of functional LTR retrotransposons by VLP DNA sequencing

Functional LTR retrotransposons form VLPs assembled from GAG proteins (Sabot and Schulman, 2006) (Supplemental Fig. S1A). Reverse transcription occurs inside the VLPs, and double-stranded cDNA products are subsequently imported into the nucleus bound to the integrase protein. After integration into new genomic loci these insertions transcribe additional gRNA. We purified VLPs after treatment with DNase I (Methods), and sequenced cDNA products from wild-type, *ddm1,* and *ddm1rdr6* using both Illumina short read and Oxford Nanopore Technologies (ONT) long read sequencing platforms (Supplemental Fig. S2). The two most productive elements, *EVADE* and the *ATCOPIA52* subfamily element AT3TE76225, constituted 1.2% and 3.4% of mapped bases, respectively, across *ddm1* short read libraries. Despite its outsized representation in the mutant libraries, AT3TE76225 has not been observed to reintegrate, and as we show later, accumulates nonproductive circular intermediates. Consequently, we named this element *SISYPHUS. EVADE* is one of two full-length elements of the *ATCOPIA93* family in *A. thaliana* Col-0, whereas the other element *ATTRAPE* is transcriptionally non-functional (Mirouze et al. 2009; Mari-Ordonez et al., 2013). When *EVADE* is transcriptionally activated, it is the most successful retroelement by far in terms of copy number increases, although transposition of elements of the *ATGP3*, *ATCOPIA13*, *ATCOPIA21,* and *ATCOPIA51* subfamilies have also been detected under non-stressed conditions (Quadrana et al., 2019; Tsukahara et al., 2009). Differential analysis of uniquely mapped VLP DNA short sequencing reads with replicates from wild-type, *ddm1* and *ddm1rdr6* mutants revealed dramatic enrichment from all of these subfamilies as well as *ATGP10*, *ATCOPIA48*, *ATCOPIA52*, and *ATCOPIA76* elements, consistent with active reverse transcription (Fig. 1; Supplemental Figs. S2,S3; Supplemental Table S1). The proportion of elements in each subfamily significantly enriched in VLP DNA-seq in each mutant are shown in Supplemental Fig. S2B. Long read coverage of *EVADE* and other active *COPIA* elements spanned the entire element and was increased in *ddm1rdr6* (Fig. 1B; Supplemental Fig. S2B; Supplemental Table S1). Furthermore, linear ecDNA was dramatically increased in *ddm1rdr6* by Southern blot (Fig. 2A). Only small numbers of linear near full-length ONT reads were found for *SISYPHUS*, *EVADE*, *ATCOPIA51* and *ATGP3* elements, suggesting the double-stranded cDNA is exported immediately after completion of reverse transcription or otherwise turned into circular DNA. In contrast, VLP DNA from *ATHILA* families were more enriched in *ddm1rdr6*, indicating regulation by easiRNA, but comprised only small fragments derived mostly from LTRs, likely reflecting abortive retrotransposition intermediates from these non-functional elements (Supplemental Fig. S3A) (Marco and Marin, 2008).

**Figure 1.**
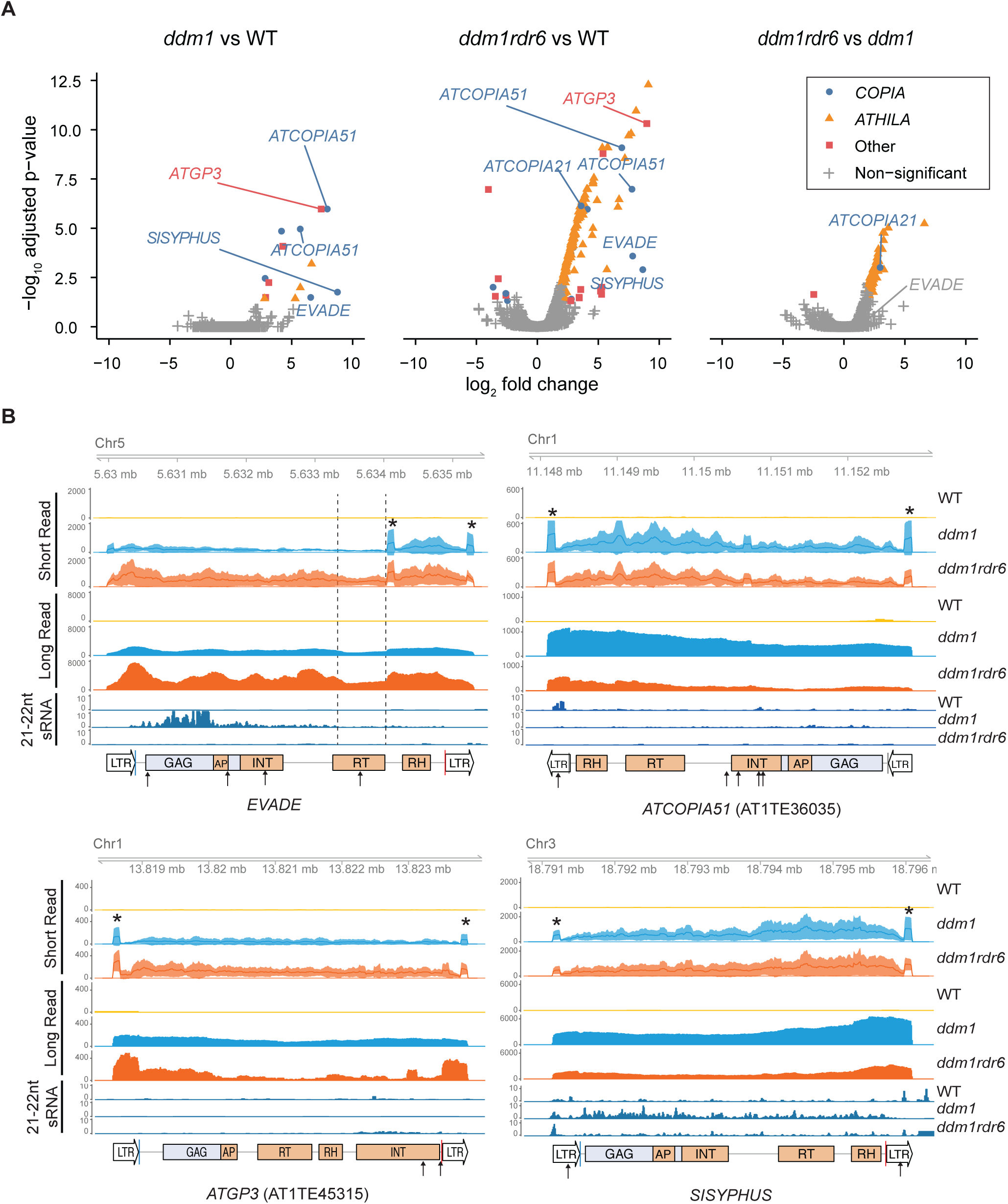
VLP DNA-seq data of LTR retrotransposons in *ddm1* and *ddm1rdr6*. (A) Differential analysis of paired-end sequencing of VLP DNA using Illumina short read platform. The statistical significance of three comparisons of wild-type (WT), *ddm1*, and *ddm1rdr6* is shown with |log_2_ (fold change)| ≥ 2 and FDR threshold at 5%. Each point corresponds to an annotated transposable element. Multiple *ATHILA* subfamilies were combined and labeled as ‘*ATHILA*’. (B) Coverage of short and long read VLP DNA-seq at representative LTR retrotransposon loci (*EVADE*, AT5TE20395; *ATGP3*, AT1TE45315; *ATCOPIA51*, AT1TE36035; *SISYPHUS*, AT3TE76225) were plotted for *ddm1* and *ddm1rdr6*. Mean read counts per million mapped reads and 95% confidence intervals of biological replicates are shown for WT (yellow, n=3), *ddm1* (blue, n=2), and *ddm1rdr6* (orange, n=3) short read libraries. VLP DNA replicate samples were pooled for each genotype and sequenced in aggregate by ONT long read sequencing. In the LTR retrotransposon annotation, abbreviations for conserved protein domains within the GAG-POL ORF are indicated as GAG, AP (amino peptidase), INT (integrase), RT (reverse transcriptase), and RH (RNase H). Blue and red lines indicate primer binding sites (PBS) and polypurine tracts (PPT). 21-22nt small RNA (sRNA) data were obtained from a previous study (Creasey et al., 2014). Target positions of miRNAs are indicated as arrows (see Supplemental Table S4 for details). Central PPT (cPPT) positions are indicated as dashed lines. Elevated coverage at the edges of strong-stop intermediate and flap DNA is shown as asterisks above *ddm1* short read data.

**Figure 2.**
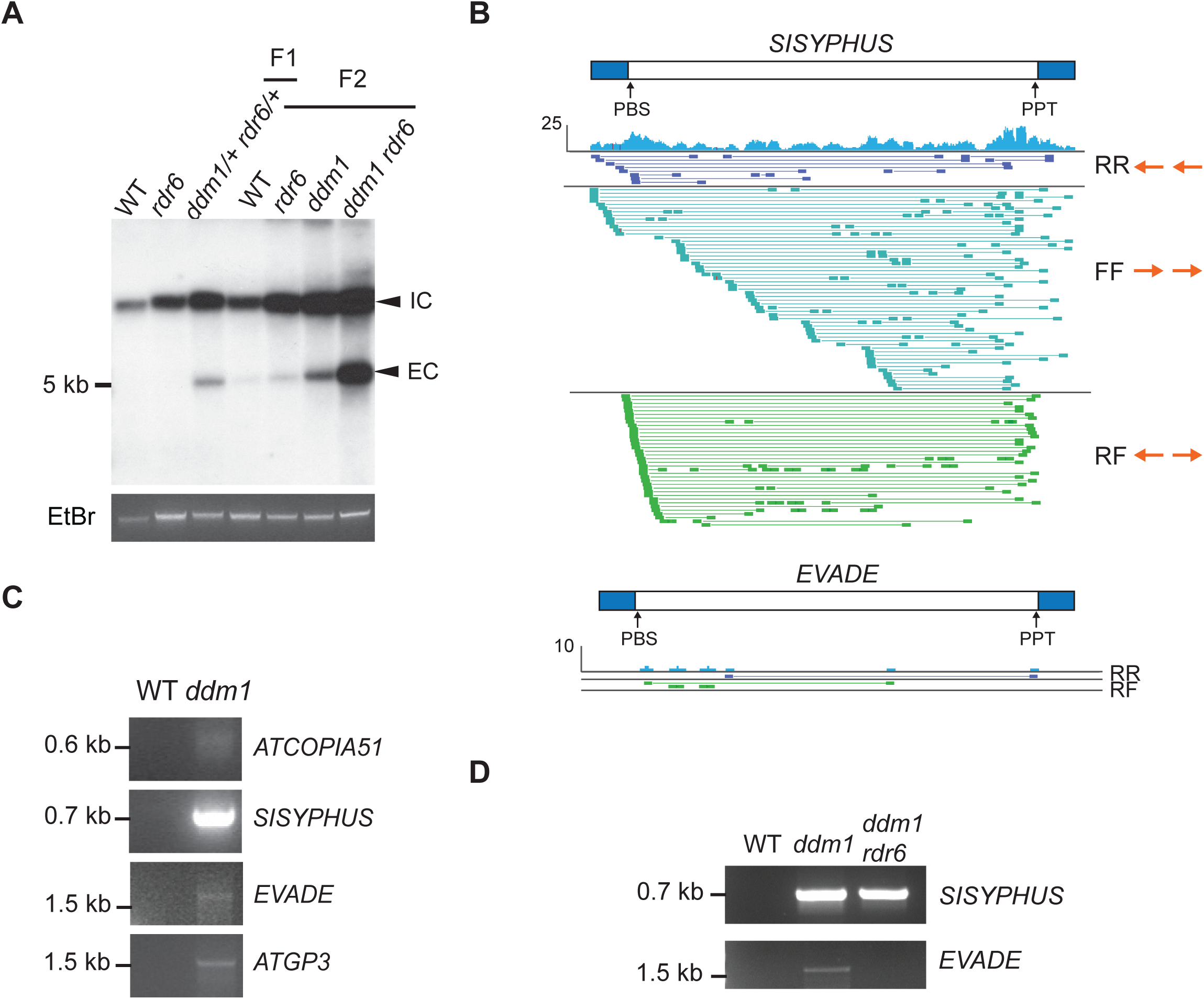
Extrachromosomal DNA of LTR retrotransposons in *ddm1* and *ddm1rdr6*. (A) Southern blotting using an *EVADE* probe was performed with undigested genomic DNA of F1 and F2 plants from the same parental lines. Integrated DNA copies (IC) and extrachromosomal DNA copies (EC) are indicated. Ethidium Bromide (EtBr) staining was used for loading control. (B) Discordant short read alignments from *SISYPHUS* (AT3TE76225) and *EVADE* in *ddm1*. Read pair orientations (forward or reverse for the first and second mate): RR and FF reads align in the same direction to the reference, indicating inversions, while RF reads face outward, indicating circular templates. LTR regions are indicated as blue bars. (C) Inverse PCR with genomic DNA to detect circular extrachromosomal DNA from *ATCOPIA51*, *SISYPHUS*, *EVADE*, and *ATGP3* in *ddm1* plants. (D) Inverse PCR with VLP DNA and reverse-forward (RF) outward reading primers for *SISYPHUS* and *EVADE*. (C-D) PCR primers are listed in Supplemental Table S6.

cDNA can exist in both linear and circular forms, and circular forms were previously reported for *EVADE* in *Arabidopsis* and a few other LTR retrotransposons in rice (Lanciano et al., 2017; Reinders et al., 2013). Outward-facing paired-end read alignments from Illumina VLP-seq reads consistent with junction-crossing reads from circular templates were absent from the vast majority of elements, but were observed in *ddm1* and *ddm1rdr6* samples for *ATCOPIA52*, *ATCOPIA93*, *ATCOPIA51*, *ATGP3* and *ATHILA* subfamily elements (Fig. 2B). *SISYPHUS* was exceptional, with over 2% of read pairs mapped non-concordantly in both mutants, whereas the proportion in *EVADE* was just 0.2% (Supplemental Table S1). Circular ecDNA formation was confirmed by inverse PCR (Supplemental Fig. S1D) whose products corresponded to one-LTR in size (Fig. 2C,D), and *SISYPHUS* was by far the most abundant. Double-stranded one-LTR circular products are thought to be generated by integrase-mediated autointegration in VLP, or as gapped intermediates in cDNA synthesis (Garfinkel et al., 2006; Munir et al., 2013; Sloan and Wainberg, 2011). In contrast, two-LTR (tandem) circular DNA with junction nucleotides is formed in the nucleus by non-homologous end joining and enhanced when integrase is non-functional (Garfinkel et al., 2006; Sloan and Wainberg, 2011). The inverse PCR products of *SISYPHUS* were one-LTR in size, suggesting the circular DNA was either a gapped double-stranded circular intermediate, or else a double-stranded product of autointegration into same strands or opposite strands (Supplemental Fig. S1), which result in deletion circles, or inversion circles, respectively (Garfinkel et al., 2006; Munir et al., 2013; Sloan and Wainberg, 2011). Both inversion and deletion circles were detected in large numbers based on outward facing reverse-forward and forward-forward paired end reads, respectively, indicating auto-integration was the major source of these circles (Fig. 2B).

In yeast, auto-integration occurs near the central PPT (cPPT) taking advantage of a DNA flap structure (Garfinkel et al., 2006). There was no strong indication of a DNA flap based on polypurine sequences and read alignment in *SISYPHUS*. We mapped individual long reads to investigate the integration sites (Fig. 3; Supplemental Figs. S1C,S3B). Deletion circles are predicted to have either the 5’ or the 3’ LTR, as well as a deleted portion of the full-length cDNA, up to the integration site, while inversion circles have an inverted portion separating the two LTR (Garfinkel et al., 2006). Strikingly, many of the *SISYPHUS* ONT reads fell into these categories, comprising either the 5’ or the 3’ LTR contiguous with a truncated or inverted portion of the retrotransposon (Supplemental Fig. S1C). These structural variants indicated the presence of circularly permuted reads, which were presumably arbitrarily sheared during library preparation. Among all the *COPIA* and *GYPSY* elements examined, only *SISYPHUS* gave rise to large numbers of these structural variants (Supplemental Table S1). The inversions spanned diverse regions of the element, consistent with inversion circles. The deleted portions terminated at inferred autointegration sites, which were distributed throughout the length of the element, consistent with the lack of a cPPT flap in *SISYPHUS*. One possibility is that nuclear import of double-stranded cDNA is not efficient for *SISYPHUS,* leading to elevated autointegration inside the VLP. This could be due to mutations in nuclear localization (Kenna et al., 1998), or else to reduced translation of the integrase gene (see below), although read distributions were comparable for *ddm1* and *ddm1rdr6*, so easiRNA likely did not play a major role.

**Figure 3.**
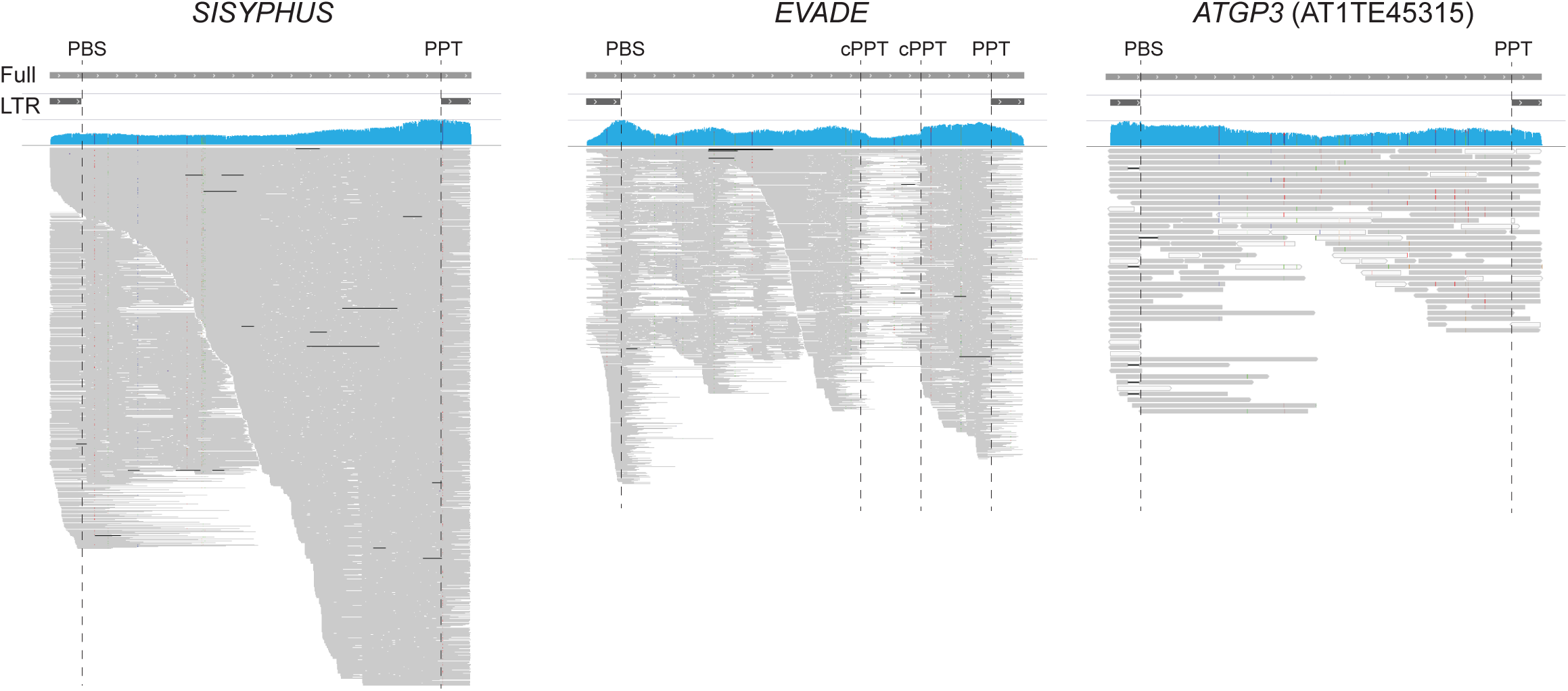
Alignments of ONT long reads from *ddm1* VLP DNA. The central polypurine tract (cPPT), PBS, and PPT positions are indicated as dashed lines relative to full and LTR annotation of *SISYPHUS* (AT3TE76225), *EVADE* (AT5TE20395), and *ATGP3* (AT1TE45315). Gaps in individual reads are indicated with black horizontal lines, and sequence mismatches are shown as colored dots in the read alignments. Pileups of linear intermediates are observed for *EVADE*, while a continuous distribution of fragment lengths is observed in *SISYPHUS*.

In sharp contrast, in *EVADE* we observed discontinuous regions of read alignments flanked by multiple cPPT, defined as 15-19 nt polypurine sequences (Figs. 1B, 3; Supplemental Fig. S3B). These regions represent active replication intermediates, generated by both minus strand and plus strand strong stop DNA, as well as extension products that terminate at cPPT and DNA flaps. The numbers of these intermediates, as well as their abundance, were significantly elevated in long-read sequencing data from *ddm1rdr6* double mutants (Figs. 1B, 3; Supplemental Fig. S3B). *ATGP3* also had elevated levels of strong stop intermediates, but few if any cPPT and no circular reads.

### 21-22nt easiRNA control retrotransposition

In a previous study, *dcl2/4* mutants lacking 21-22nt small RNA were shown to accumulate 24nt small RNA from an *EVADE* transgene driven by an ectopic promoter, leading to transcriptional silencing (Mari-Ordonez et al., 2013). In contrast, *rdr6* had no effect on the *EVADE* transgene, which can be interpreted as evidence that easiRNA might not inhibit transposition in wild-type cells. We tested whether easiRNA contribute to control of endogenous *EVADE* in *ddm1* and *ddm1rdr6* mutants by analyzing DNA copy numbers and RNA levels using qPCR and RT-qPCR (Methods). Both *ddm1* and *ddm1rdr6* contained higher copy numbers of *EVADE* than wild-type implying high rates of transposition, while copy numbers of *ATGP3* and *SISYPHUS* remained constant. Using quantitative PCR, we detected an increase from 2 copies of *EVADE* in wild-type to 12 copies in *ddm1* to 40 copies in *ddm1rdr6* F2 siblings (Fig. 4A). Similar increases were observed in F2 and F3 *rdr6* progeny from a parental *+/rdr6* F1 with active *EVADE* elements (Fig. 4C) inherited epigenetically (Mari-Ordonez et al., 2013). We detected parallel increases in gRNA levels reflecting these increases in copy number (Fig. 4B,D). Consistent with gRNA levels, extrachromosomal *EVADE* copies were also more abundant in *ddm1rdr6* than in *ddm1* (Fig. 2A). RNase H cleavage products just upstream of the PBS, which are a hallmark of active transposition (Schorn et al., 2017), were readily detected for *EVADE* in both *ddm1* and *ddm1rdr6* (Supplemental Fig. S4A,B). We conclude that easiRNA actually inhibit *EVADE* retrotransposition, in *ddm1* mutants by downregulating RNA levels.

**Figure 4.**
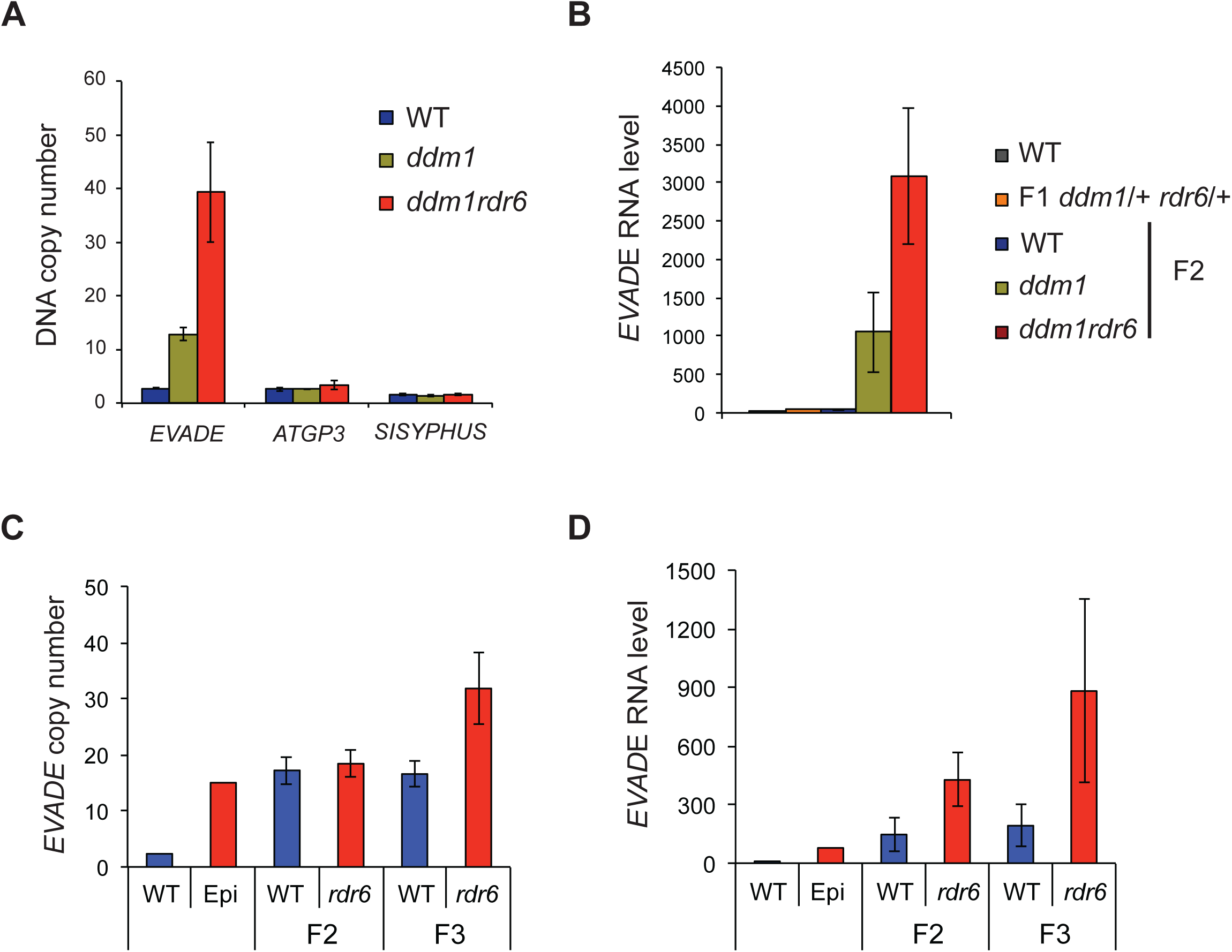
DNA and RNA levels of LTR retrotransposons in *ddm1* and *rdr6* mutants. (A) DNA copy numbers of *EVADE*, *ATGP3*, and *SISYPHUS* in *ddm1* and *ddm1rdr6* were normalized with a single copy gene (AT5G13440). (B) RT-qPCR data of *EVADE* elements using POL primers. Y-axis indicates relative levels of *EVADE* genomic RNA to wild-type (WT) after normalization to *ACT2*. (C-D) *EVADE* DNA copy number and genomic RNA levels were analyzed in F2 and F3 progenies of F1 plants carrying active *EVADE* epigenetically inherited from parental *rdr6*/+ (Epi) crossed with WT pollen. Error bars indicate standard deviations (n=3).

In backcrosses to wild-type (WT) plants, *EVADE* activity is inherited epigenetically but copy number increases are thought to be limited by a switch from 21nt to 24nt siRNA, accompanied by re-methylation and silencing (Mari-Ordonez et al., 2013; Mirouze et al., 2009; Reinders et al., 2013). Interestingly, active *EVADE* elements can be re-silenced through the female gametophyte, but not through the male gametophyte (Reinders et al., 2013) where easiRNA normally accumulate (Borges et al., 2018; Slotkin et al., 2009). We sequenced small RNA from wild type and *ddm1* flower buds and pollen, and found that 21-22nt easiRNA from *EVADE* were abundant in *ddm1* inflorescence tissues, but absent from pollen (Fig. 5). In contrast, *ATHILA2* and *ATHILA6A* easiRNA were present in wild type pollen (Slotkin et al., 2009), while *ATCOPIA31* 21-22nt easiRNA were strongly upregulated in *ddm1* pollen. Thus the absence of *EVADE* easiRNA in pollen must be due to transcriptional repression independent of DDM1, and likely accounts for the lack of paternal re-silencing (Reinders et al., 2013).

**Figure 5.**
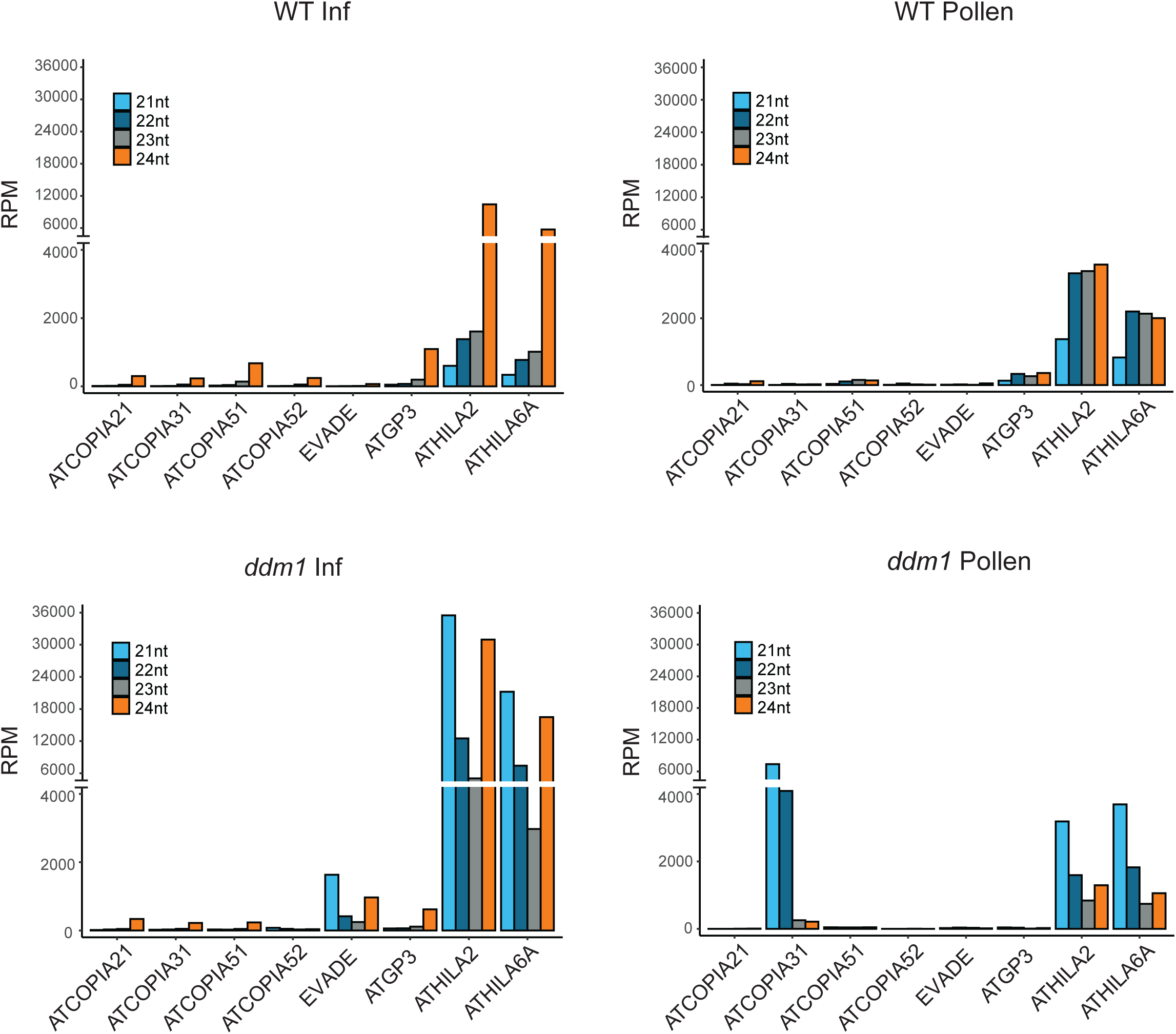
Small RNA profiles of representative LTR retrotransposons. 21, 22, 23, and 24nt small RNA levels in inflorescence tissues and pollen of wild-type (WT) and *ddm1*. Reads per million (RPM) was calculated from entire elements including LTR and coding sequences.

### Post-transcriptional suppression by easiRNA

Since easiRNA in *ddm1* mutants depend on AGO1 associated with 21-22nt small RNA (Nuthikattu et al., 2013), and AGO1 represses translation of target mRNA (Li et al., 2013), we tested whether easiRNA can affect translation efficiency of transposon transcripts. Translating ribosome affinity immunopurification (TRAP) RNA-seq has been utilized to estimate polysomal occupancy and translation efficiency in plants (Juntawong et al., 2014). Furthermore, microsome-polysomal fractionation has revealed that microRNA-dependent translational control takes place on the endoplasmic reticulum (Li et al., 2013). We generated TRAP lines of *35S:FLAG-RPL18* in *ddm1* and *ddm1rdr6* mutant backgrounds, and performed total RNA-seq, total-polysomal RNA-seq, and microsome-polysomal RNA-seq. The polysomal RNA occupancy (Polysomal RNA / Total RNA) was obtained for 3903 transposable elements defined as open reading frames from TAIR10 annotation (see Methods). As for the comparison between *ddm1* and *ddm1rdr6*, we could detect the effect of the *rdr6* mutation in microsome-polysomal RNA-seq data for known targets of RDR6, such as *ARF4* (Marin et al., 2010), and for a handful of transposons (Fig. 6A; Supplemental Fig. S5; Supplemental Tables S2,S3). Among 31 up-regulated transposons in *ddm1rdr6* relative to *ddm1*, 26 elements belonged to *ATHILA* LTR retrotransposon families (Supplemental Table S3), which are a major source of RDR6-dependent easiRNA. Although *ATHILA* elements in *A. thaliana* cannot transpose, a subgenomic mRNA encoding ORF2 (the “env” gene) is spliced from the full-length mRNA (Havecker et al., 2004; Wright and Voytas, 2002), and was enriched on polysomes (Supplemental Fig. S5; Supplemental Table S3). This subgenomic RNA is targeted extensively by miRNA which trigger easiRNA production (Creasey et al., 2014). Interestingly, the other 3 elements were *ATENSPM3*, *LINE1_6* and *VANDAL3*, all of which have been identified as active elements in *ddm1* mutants, or in population level studies of transposon variants (Stuart et al., 2016). These non-LTR and DNA transposons are also targets of miRNA and generate RDR6-dependent easiRNA (Creasey et al., 2014). *EVADE* easiRNA are generated from the GAG subgenomic RNA (Mari-Ordonez et al., 2013), but polysomal occupancy was not increased in *ddm1rdr6* (Fig. 6B). GAG subgenomic mRNA from *SISYPHUS* was highly enriched in polysomes, consistent with previous studies (Oberlin et al., 2017), whereas the relative abundance of *EVADE* POL transcripts on polysomes indicates higher translation rates of integrase and reverse transcriptase (Oberlin et al., 2017). Unlike for *ATHILA*, polysome association of *COPIA* transcripts was unaffected by RDR6.

**Figure 6.**
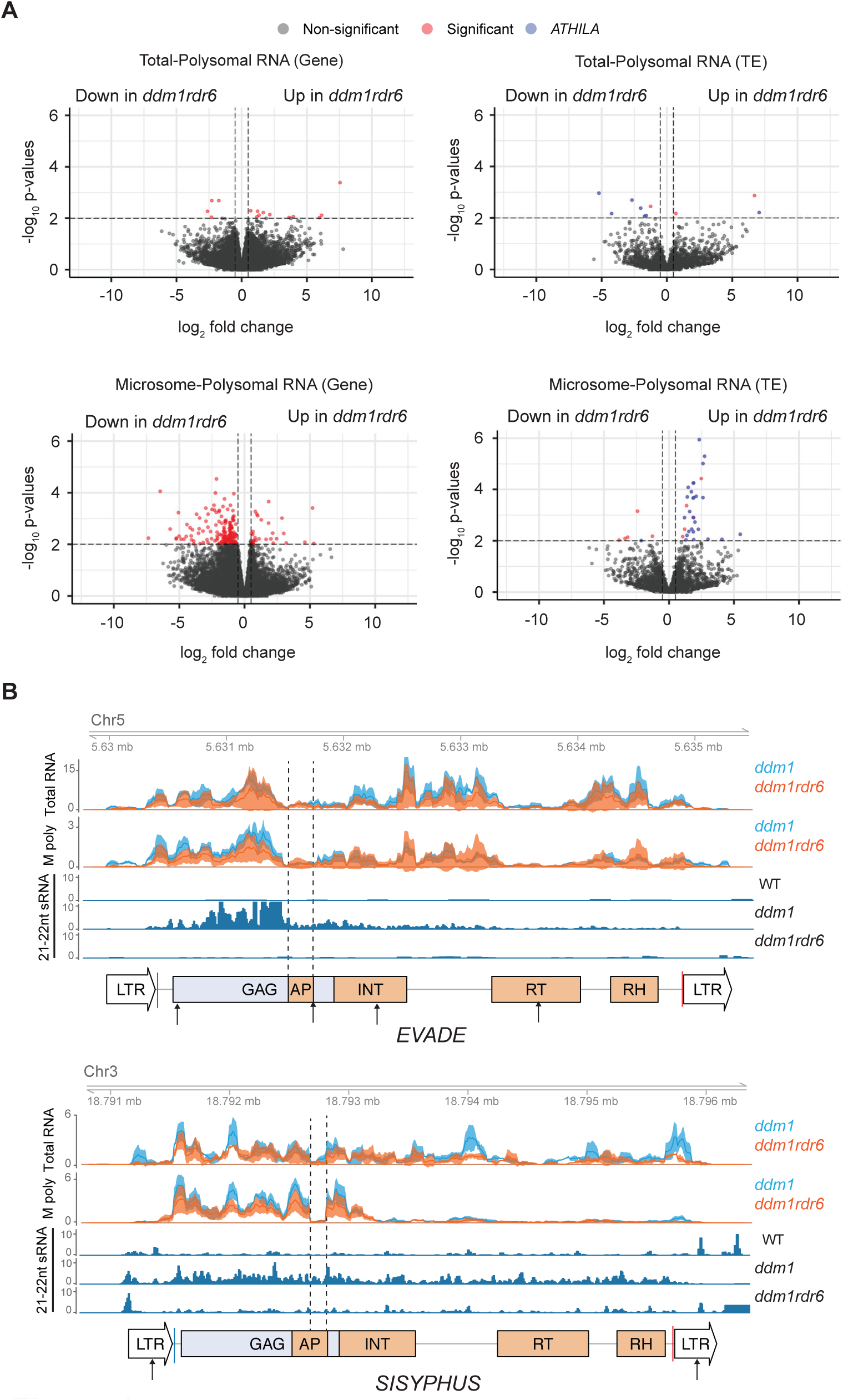
Translatome profiles of *ddm1* and *ddm1rdr6*. (A) Differential analysis of polysomal RNA-seq data between *ddm1* and *ddm1rdr6*. Polysomal RNA-seq values were normalized by total RNA seq values to reflect polysomal enrichment (Methods). Red dots indicate significantly regulated genes or transposable elements (TE) by cut-off values of |log_2_ (fold change)| > 0.5 and p-values < 0.01 which include *ARF4* as an internal control. Significantly regulated *ATHILA* family elements are labeled with blue dots. (B) Total RNA and microsome-polysomal RNA (M poly) levels are shown for *EVADE* (AT5TE20395) and *SISYPHUS* (AT3TE76225). Mean read counts per million mapped reads and 95% confidence intervals of three biological replicates are shown for *ddm1* (blue) and *ddm1rdr6* (orange). Conserved protein domains, PBS and PPT, small RNA profiles and miRNA target sites are indicated as in Fig. 1.

easiRNA require miRNA triggers that target these transcripts (Creasey et al., 2014), and *SISYPHUS* LTRs were targeted by a single miRNA in the R region of the LTR. Consistent with this miRNA acting as a trigger, easiRNA accumulated along the length of the mRNA between the LTRs (Fig. 6B; Supplemental Table S4). In the case of *EVADE*, 4 miRNA were predicted to target the gRNA somewhere along its length. Remarkably, miR2938 was predicted to target the start codon of the GAG gene immediately 5’ of the easiRNA cluster, while miR5648-5p targets the 3’ end of the easiRNA cluster (Supplemental Fig. S4C,D; Supplemental Table S4). *EVADE* easiRNAs were also down-regulated in *ddm1dcl1* as compared to *ddm1* (Supplemental Fig. S4E) suggesting that miRNA were involved (Creasey et al., 2014). miR2938 and miR5648-5p expression were reported in pollen and root cells (Breakfield et al., 2012; Grant-Downton et al., 2009). We did not detect miRNA-mediated cleavage by PARE-seq (Creasey et al., 2014) or by RACE-PCR in inflorescence tissues, but secondary siRNA, such as easiRNA, do not require cleavage so long as miRNA recognition recruits RdRP (Axtell et al., 2006; de Felippes et al., 2017). Consistent with induction without cleavage, *EVADE* easiRNA were not phased (Arribas-Hernandez et al., 2016). miR5663 was detected in inflorescence tissues (Supplemental Fig. S4F), and targets the *EVADE* intron near the splice acceptor site (Supplemental Fig. S4C). Interestingly, the level of unspliced RNA was increased in *ddm1dcl1* mutants (Supplemental Fig. S4G), indicating that miR5663 might target unspliced gRNA and promote the accumulation of spliced GAG RNA, but further experiments would be required to demonstrate this requirement. Negative regulation of *P*-element splicing by piRNA has been reported in Drosophila (Teixeira et al. 2017). *ATCOPIA21* and *ATCOPIA51* had no strongly predicted miRNA targets, and easiRNA were barely detected in somatic tissues (Fig. 1B; Supplemental Fig. S3A) (Oberlin et al., 2017) accounting for lack of regulation by *RDR6*. In contrast, significant levels were detected in pollen (Fig. 5) (Borges et al., 2018), where most gypsy and copia class retrotransposons are targeted by miR845, a pollen specific miRNA that targets the primer binding site (Borges et al., 2018).

### Transcriptional repression by easiRNA

In plants, both 21/22nt easiRNA and 24nt siRNA species have the capacity to direct DNA methylation via the RNA directed DNA methylation (RdDM) pathway (Borges and Martienssen, 2015). As such, we sought to more comprehensively define the impact of RDR6-dependent easiRNA on transposable elements at the transcriptional level. Genome-wide analysis of bisulphite sequencing revealed only minimal differences in DNA methylation between *ddm1* and *ddm1rdr6* (Creasey et al., 2014). However, transcriptional repression can also be achieved via repressive histone modifications, such as histone H3 lysine-9 dimethylation (H3K9me2) which, in many organisms, is known to be guided by small RNA (Fagegaltier et al., 2009; Gu et al., 2012; Martienssen and Moazed, 2015; Volpe et al., 2002). We therefore performed H3K9me2 ChIP-seq in wild-type, *ddm1* and *ddm1rdr6* plants. As expected, *ddm1* mutants showed genome-wide loss of H3K9me2 relative to wild-type (Supplemental Fig. S6; Supplemental Table S5). We identified a subset of regions that, in contrast to the rest of the genome, maintained high levels of H3K9me2 in *ddm1*. These loci were composed almost entirely of *ATHILA* family elements and lost H3K9me2 in *ddm1rdr6* mutants (Fig. 7; Supplemental Fig. S6). The RDR6-dependent accumulation of H3K9me2 at these loci was significantly correlated with levels of 21-22nt easiRNA accumulation (Fischer’s, p=3e-10), and the subset of *ATHILA* elements regulated by RDR6-dependant H3K9me2 had higher levels of 21-22nt and 24nt sRNAs than those that were unaffected (Supplemental Fig. S6C). In contrast, *COPIA* elements were not associated with RDR6-dependent H3K9me2 (Fischer’s, p=0.98), showing only modest increases in H3K9me2 and 24nt siRNAs in *ddm1rdr6* (Supplemental Fig. S7). Taken together, these results strongly imply an as yet unexplored role for easiRNA in transcriptional control of transposable elements via targeting of the repressive histone modification H3K9me2.

**Figure 7.**
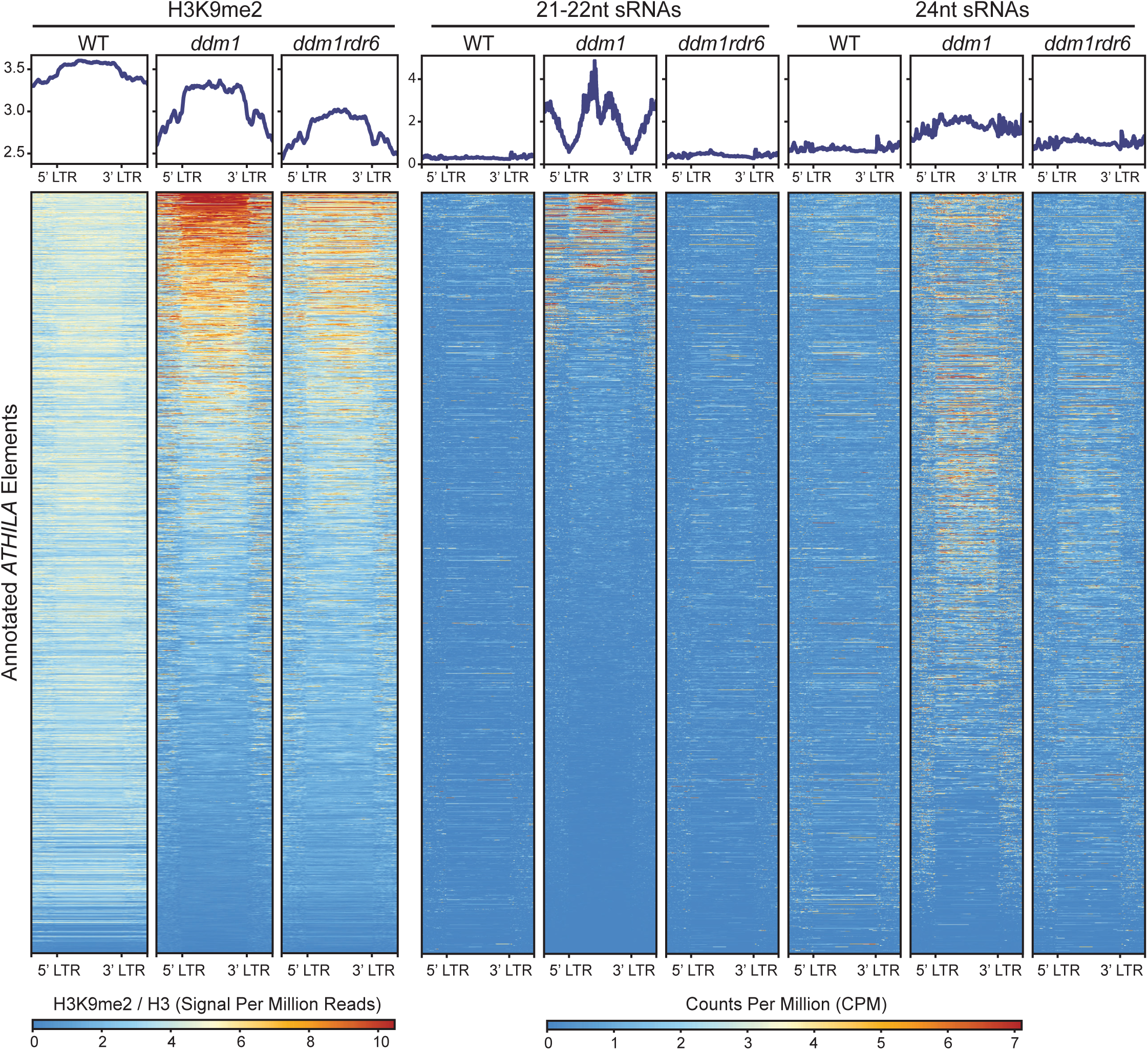
*ATHILA* family elements gain RDR6-dependent H3K9me2 in *ddm1*. H3K9me2 signal at transposable elements from multiple *ATHILA* families was analyzed in wild-type (WT), *ddm1*, and *ddm1rdr6* genotypes and correlated with previously published small RNA data (Creasey et al., 2014). RDR6-dependent gains in H3K9me2 co-localize with increased 21-22nt siRNAs in *ddm1*. Plots depict transposable elements annotations scaled to 5kb, as well as 2kb upstream and downstream of each feature. H3K9me2 ChIP data was normalized by H3, and small RNA data was normalized by counts per million.

## Discussion

Sequencing of VLP DNA detected all known functional LTR retrotransposons in *Arabidopsis*, as well as some non-functional ones. Full-length VLP DNA read coverage from *ATCOPIA* and *ATGP* families (Fig. 1B; Supplemental Fig. S3A) corresponded to relatively young and low-copy elements known to transpose. Ancient *ATHILA* elements did not make full-length VLP DNA confirming these gypsy retrotransposons are non-functional (Havecker et al., 2004; Marco and Marin, 2008), but short products matching the LTR appeared to correspond to aborted strong stop replication intermediates (Supplemental Fig. S3A). Interestingly, similar LTR fragments from *ATHILA2* comprise an important family of dispersed centromeric satellite repeats known as 106B (May et al., 2005; Thompson et al., 1996), and retrotransposition might account for their origin. Thus functional and non-functional retrotransposons could be readily distinguished even though non-functional *ATHILA* elements are present in copy numbers 3 to 4 orders of magnitude higher than functional *ATCOPIA* and *ATGP* elements. As for *SISYPHUS*, non-productive one-LTR circular DNA, corresponding to autointegration “suicide” products, markedly accumulated in the VLP at levels far higher than productive retrotransposons such as *EVADE*. In contrast, two-LTR circular products of *SISYPHUS* were very rare, whereas small amounts of *EVADE* two-LTR products were present as previously described (Reinders et al., 2013), presumably due to recombination of non-integrated copies by host DNA repair enzymes in the nucleus. Both short read and long read sequencing revealed that these auto-integration products in *SISYPHUS* VLP led to non-functional deletion and inversion circles, accounting for lack of transposition.

We found that some retrotransposons are regulated post-transcriptionally by RNA interference, while others are regulated at the transcriptional level by histone H3 lysine-9 methylation guided by small RNA. RDR6-dependent easiRNA inhibit retrotransposition via post-transcriptional silencing of genomic RNA, by translational suppression of subgenomic RNA, and by controlling transcription via histone modification. *ATHILA* elements are no longer functional, but they are the primary source of easiRNA which arise by miRNA targeting of a spliced subgenomic RNA encoding the “ENV” protein (Creasey et al., 2014). These easiRNA inhibit polysome association of this subgenomic RNA, and also inhibit transcript levels by guiding histone H3K9me2. This transcriptional silencing occurred in the absence of DNA methylation in *ddm1* mutants. In plants, RNAi dependent histone modification is thought to depend on RNA dependent DNA methylation, found in asymmetric CHH contexts. As CHH methylation stays more or less the same in *ddm1*, while H3K9me2 is increased (Fig. 7), this might indicate the existence of a novel pathway for RNA guided histone methylation, resembling that found in Drosophila, *C.elegans* and fission yeast, which lack DNA methylation. Further investigation will be required to establish if such a pathway exists.

In contrast to *ATHILA*, linear extrachromosomal copies of *EVADE* accumulated in *ddm1* and were further enriched by mutations in *RDR6*. Like *ATHILA*, *EVADE* is targeted by 3 or 4 miRNA that likely trigger easiRNA from the subgenomic GAG gene transcript, which is found associated with polysomes (Oberlin et al., 2017). However, association of the *EVADE* GAG mRNA with polysomes was unaffected in *ddm1rdr6* mutants. Instead, levels of gRNA increased 3-fold, suggesting that *EVADE* easiRNA act postranscriptionally to target gRNA directly. *SISYPHUS* easiRNA arose from full-length gRNA between the two LTR. Polysomal association of full-length *EVADE* GAG-POL is far more abundant than *SISYPHUS* GAG-POL, although both were unchanged in the absence of *RDR6* (Fig. 6). As the integrase protein is translated from this transcript, this could contribute to lack of nuclear integration of *SISYPHUS* relative to *EVADE*. Thus, while easiRNA have a significant impact on *COPIA* gRNA accumulation, and so inhibit increases in copy number, they have only limited impact on translation. 22nt tRNA-derived small RNA fragments (3’CCA-tRFs) were recently shown to inhibit endogenous retroviruses (ERV) in mammalian cells by targeting the PBS by RNA interference (Schorn et al., 2017), and it is possible that *EVADE* easiRNA may have a similar function in plants.

In conclusion, long read and short read sequencing of VLP DNA has revealed features that distinguish functional and non-functional replication intermediates, and provides a powerful tool for identifying active transposons from complex genomes, and for investigating molecular signatures of LTR retrotransposons. One such feature is the cPPT, which is present in *EVADE* but absent in *SISYPHUS*. cPPT are hallmarks of the most active retrotransposons including Ty1 in yeast, as well as HIV and other lentiviruses, where cPPT are thought to be important for nuclear import of double-stranded cDNA (VandenDriessche et al., 2002; Zennou et al., 2000). Our work shows that these features may play a significant role in the activity of *EVADE*, the most active retrotransposon in *Arabidopsis*, and that their absence may account for the lack of nuclear integration of *SISYPHUS*, and high levels of “suicide” by autointegration. By comparing VLP sequencing, transcriptome sequencing and translatome sequencing we have been able to establish the multiple levels at which easiRNA regulate the *Arabidopsis* LTR retrotransposons. Our methods are widely applicable to other plant and animal models and to human cells, especially those with genomes that contain very large numbers of non-functional LTR retrotransposons. Leveraging long read sequencing, complete sequences of active transposon intermediates can be studied even when no reference genome is available.

## Methods

### Plant materials

All genotypes in this study are Col-0 background including wild-type, *dcl1-11*, *ddm1-2*, and *rdr6-15.* Genotyping primers are listed in Supplemental Table S6. Homozygous plants of *ddm1-2* and *ddm1-2 rdr6-15* were generated from heterozygous *ddm1-2* backcrossed five times with Col-0 (*ddm1-2* BC5), and their 2^nd^ generation was used for VLP DNA-seq experiments. For polysomal RNA-seq experiments, inbred *ddm1-2* was independently crossed to *35S:FLAG-RPL18* and to *rdr6-15 35S:FLAG-RPL18*. The F3 plants were used for polysomal RNA purification.

### gDNA extraction and DNA analyses

Whole inflorescence stems of 4 week-old *Arabidopsis* plants were frozen and ground in liquid nitrogen. Total gDNA was isolated using Nucleon PhytoPure kit (GE healthcare). *EVADE* DNA copy number was quantified using qPCR with *EVADE* qPCR primers and single copy gene primers as reference (the primers are listed in Supplemental Table S6). Southern blotting was performed using *EVADE* Probe B as described (Mirouze et al., 2009).

### Chromatin immunoprecipitation (ChIP)

ChIP was performed with two biological replicates of 10-d-old seedlings using H3K9me2 (Abcam; ab1220) and H3 (Abcam; ab1791) antibodies, following a previously described protocol (Ingouff et al., 2017). Sequencing libraries were prepared using NEBNext Ultra II DNA Library Prep Kit for Illumina (New England Biolabs) with size selection for ~200 bp insert DNA. The ChIP-seq libraries were sequenced using Illumina NextSeq High Output SR 76 with 76-cycle single reads. Two biological replicates were prepared and sequenced for each genotype of interest. Prior to alignment, adapter trimming was performed using Trimmomatic (Bolger et al., 2014) and read quality was assessed with FastQC (http://www.bioinformatics.babraham.ac.uk/projects/fastqc). Reads were aligned to the TAIR10 reference genome using BWA-MEM (Li, 2013) with default parameters. Only primary alignments were retained, and optical and PCR duplicates were removed using SAMtools (Li et al., 2009). Peak calling was performed using MACS2 (Zhang et al., 2008) broad peak calling with a q-value cutoff of 0.05 and normalization by signal per million reads. Peaks that were differentially regulated across genotypes were identified using MAnorm (Shao et al., 2012) and confirmed between biological replicates. Annotation of these differentially regulated peaks was performed using a combination of BEDOPS (Neph et al., 2012) tools and custom scripts. deepTools (Ramirez et al., 2014) was used to visualize the data.

### RNA extraction and RT-qPCR

Total RNA was isolated from the same tissues used for gDNA extraction with Direct-zol RNA MiniPrep Plus (Zymo Research). DNase I was treated on column. cDNA was synthesized with SuperScript VILO Master Mix (Thermo Fisher Scientific). qPCR was performed using iQ SYBR Green Supermix. Primers are listed in Supplemental Table S6.

### Polysomal RNA-seq

Total polysome was isolated using ribosome immunopurification as described previously (Mustroph et al., 2009; Mustroph et al., 2013). Briefly, inflorescence tissues of *FLAG-RPL18* lines were ground in liquid nitrogen and transferred to polysome extraction buffer (PEB). Cell debris was removed by centrifugation and filtering with miracloth. The supernatant was taken and transferred to pre-washed EZview anti-FLAG agarose beads (Sigma) for 2 h at 4 °C. The agarose beads containing polysomes were washed once with PEB and three times with washing buffer. Polysomes were eluted using 3X FLAG peptide (Sigma) and used for RNA extraction with Direct-zol RNA miniprep kit (Zymo Research) including DNase I treatment. Ribosomal RNA (rRNA) in the samples was depleted by Ribo-Zero Magnetic Kit (Plant Leaf) (Epicentre). Then, rRNA free samples were used for RNA-seq library preparation using ScriptSeq v2 RNA-Seq Library Preparation Kit (EPicentre). Microsome-polysomal RNA was obtained using a previously described method with some modifications (Li et al., 2013). Briefly, 2 g frozen tissues were suspended to 7 ml microsome extraction buffer (MEB). After removing cell debris by filtration with micracloth and centrifugation at 10,000g for 15 min at 4°C, the supernatant was transferred on the top of 1.7M/0.6M sucrose cushions and applied to ultracentrifugation using swing rotor at 140,000g for 1 h at 4°C. The microsome fraction of the 1.7M/0.6M layer interface was harvested and diluted 10 times by MEB and centrifuged at 140,000g for 0.5 h at 4 °C to obtain microsome pellet. The pellet was re-suspended with 8 ml PEB and used for ribosome immunopurification and RNA-seq library preparation as described above. The PE 101 sequencing data was obtained using Illumina HiSeq 2000 platform. The paired-end reads were mapped to *Arabidopsis* TAIR10 genome using TopHat and the polysome occupancy (Polysomal RNA / Total RNA) was calculated using systemPipeR package (Backman and Girke, 2016) with raw count data obtained by Cuffnorm.

### VLP DNA-seq

Virus-like-particles were purified using modified method reported previously (Bachmair et al., 2004). 4 g of 4 week-old whole inflorescence stems were ground with 10 ml of ice-cold VLP extraction buffer and 10 ml of sea sand on ice. 10 ml of the extraction buffer and Triton X-100 were added and mixed. The slurry was transferred to a 50 ml tube and centrifuged for 5 min at 180g and 4 °C. The supernatant was carefully transferred onto 5 ml of prechilled 15% sucrose, 10 mM potassium phosphate buffer, pH 7.2 and ultracentrifuged for 1.5 h at 109,000g and 4 °C using fixed angle rotor. The pellet was washed with the 15% sucrose buffer and resuspended with 4 ml particle suspension buffer to obtain VLP fractions. To remove non-VLP DNA, 0.5 ml of the VLP sample was treated with 5 μl of 1 mg/ml DNase I at 37°C for 10 min. 20 μl of 0.25 M EDTA, 50 μl of 10% SDS, 25 μl of 10 mg/ml proteinase were added and incubated at 65°C for 10 min. VLP DNA was purified by 0.5 ml equilibrated (pH 8.0) phenol:chloroform:IAA (25:24:1) mixture three times and with 0.5 ml chloroform:IAA (24:1) once. The last aqueous fraction was transferred into a new 1.5ml tube and used for 100% ethanol precipitation with 40 μl 3M sodium acetate, pH 7.0. The DNA pellet was washed with 70% ethanol, dried, and resuspended with 100 μl TE buffer. 1 μl of RNase A (10 mg/ml) was added to the VLP DNA sample and incubated 10 min. The treated DNA sample was purified using DNA Clean & Concentrator (Zymo Research). The DNA was sheared to 650 bp using Covaris S220 and subsequently used for DNA-seq library preparation with NEBNext Ultra DNA Library Prep Kit (New England Biolabs). The paired-end sequencing datasets with 101 nt read length (PE101) were obtained by Illumina HiSeq 2000. Adapters were trimmed from raw reads with Skewer (Jiang et al., 2014) in paired-end mode and read pairs with both mates longer than 25 nt were retained. Reads were aligned to the TAIR10 genome with STAR (Dobin et al., 2013) in two-pass mode to improve spliced alignment at unannotated introns. Intact bacteria co-purified with VLP, as indicated by large numbers of reads mapping to bacterial genomes (up to 95% in WT), and these were discarded. Reads mapping equally well to multiple locations were randomly assigned, and chimeric/split read alignments were output separately from concordant alignments. Optical and PCR duplicates were removed from the alignments with the Picard toolkit (http://broadinstitute.github.io/picard). Counts of reads mapping to the TAIR10 transposon annotations were computed with featureCounts (Liao et al., 2014). Pairwise differential expression at TAIR10 transposon loci was tested across three wild-type, two *ddm1*, and three *ddm1rdr6* replicates using quasi-likelihood F-tests in edgeR (Robinson et al., 2010), controlling FDR at 5% and a log_2_(fold-change) threshold of 2.

Oxford Nanopore Technologies long-read libraries were prepared as follows: 10 ng per genotype of purified VLP DNA extract was pooled from the replicate samples and initially amplified following the conditions in the “1D Low-input genomic DNA with PCR” (SQK-LSK108) protocol with reagents. End-repair, dA-tailing and PCR adapter ligation were performed, followed by 16 cycles of PCR amplification. PCR products were purified and concentrated with Ampure XP beads (Agencourt), and 300 ng of eluate per sample was carried through to library preparation following the “1D Genomic DNA by Ligation” protocol with SKQ-LSK109 reagents. Libraries were loaded onto r9.4 (FLO-MIN106) flow cells and sequenced on a GridION X5. Basecalling was performed offline with Guppy v2.3.1 using the default r9.4.1 model. Using Porechop (https://github.com/rrwick/Porechop), ONT sequencing adapters were trimmed from 5’ ends, 3’ ends, and the middle of raw reads. Reads with middle adapters were split. Remaining reads longer than 100 bp were aligned to the TAIR10 reference with Minimap2 (Li, 2018) for coverage and read alignment plots. Structural variants were called on NGMLR (Sedlazeck et al., 2018) alignments using Sniffles (Sedlazeck et al., 2018) with default parameters, except minimum read support was reduced to 3. Ribbon was used to visualize complex structural variants (Nattestad et al., 2016).

### 5’ RACE PCR

5’ RACE PCR was performed using FirstChoice RLM-RACE Kit (Thermo Fisher Scientific) without the treatments of calf intestine alkaline phosphatase and tobacco acid pyrophosphatase. A gene-specific primer was used for cDNA synthesis after adaptor ligation (Supplemental Table S6). 1^st^ and 2^nd^ nested PCR was performed with the primers are listed.

### Small RNA-seq data

Small RNA-seq libraries from inflorescence and pollen for comparisons of 21, 22, and 24nt small RNA between wild-type and *ddm1* were prepared as previously described (Borges et al., 2018). Wild-type pollen sample was previously deposited in the Gene Expression Omnibus (GEO) database (GSM2829912). Briefly, small RNAs were purified by running total RNA from pollen and inflorescence tissues on acrylamide gels (15% polyacrylamide, 7 M urea) with size-selection of 18-to-30-nt regions. Small RNAs were extracted from the gel bands using Trizol LS (Life Technologies) and Direct-zol columns (Zymo Research). Libraries were prepared with the TruSeq small RNA sample preparation kit (Illumina) and sequenced in Illumina MiSeq platform. Data analysis was done as previously reported (Borges et al., 2018). 21-22nt small RNA datasets from inflorescence (Creasey et al., 2014) were obtained from NCBI GEO accession GSE52951. After adapter trimming with Skewer, reads were quality filtered with fastp (Chen et al., 2018) and aligned to the TAIR10 genome with ShortStack (Axtell, 2013) with default parameters except “--bowtie_m 1000 --ranmax 50”.

### LTR Retrotransposon Annotation

GenomeTools was used to structurally annotate retrotransposons across the TAIR10 genome. First, LTRharvest (Ellinghaus et al., 2008) was run to detect LTR sequences with at least 85% similarity separated by 1-15 kbp flanked by target site duplications and the TGCA motif. Then, LTRdigest (Steinbiss et al., 2009) was run to annotate internal transposon features including the PBS, PPT, and GAG and POL protein homology.

### Genome Browser Figures

Genome-wide read coverage for VLP DNA, small RNA, total and polysomal RNA libraries was calculated with bamCoverage from deepTools (Ramirez et al., 2014) and normalized to reads per nucleotide per million mapped reads and plotted across the genome with Gviz (Hahne and Ivanek, 2016) or IGV (Thorvaldsdottir et al., 2013).

## Data Access

All raw and processed sequencing data generated in this study have been submitted to the NCBI Gene Expression Omnibus (GEO; https://www.ncbi.nlm.nih.gov/geo/) under accession number GSE128932.

## Acknowledgements

We thank Vincent Colot, Leandro Quadrano, Tetsuji Kakutani, Matthias Benoit, and all members of the Martienssen laboratory for discussions. Research in the Martienssen laboratory is supported by the US National Institutes of Health (NIH) grant R01 GM067014, The National Science Foundation Plant Genome Research Program, and by the Howard Hughes Medical Institute. The authors acknowledge assistance from the Cold Spring Harbor Laboratory Shared Resources, which are funded in part by the Cancer Center Support Grant (5PP30CA045508).

## Authors’ contributions

SCL, EE and RM designed the study; SCL, EE, FB, JSP, and PL performed the experiments; SCL, EE, BB, FB, and AS analyzed the data and its significance; SCL, EE, BB, and RM wrote the manuscript.

