## Supplemental Fig. S1 for "*Arabidopsis* LTR retrotransposons and their regulation by epigenetically activated small RNA"

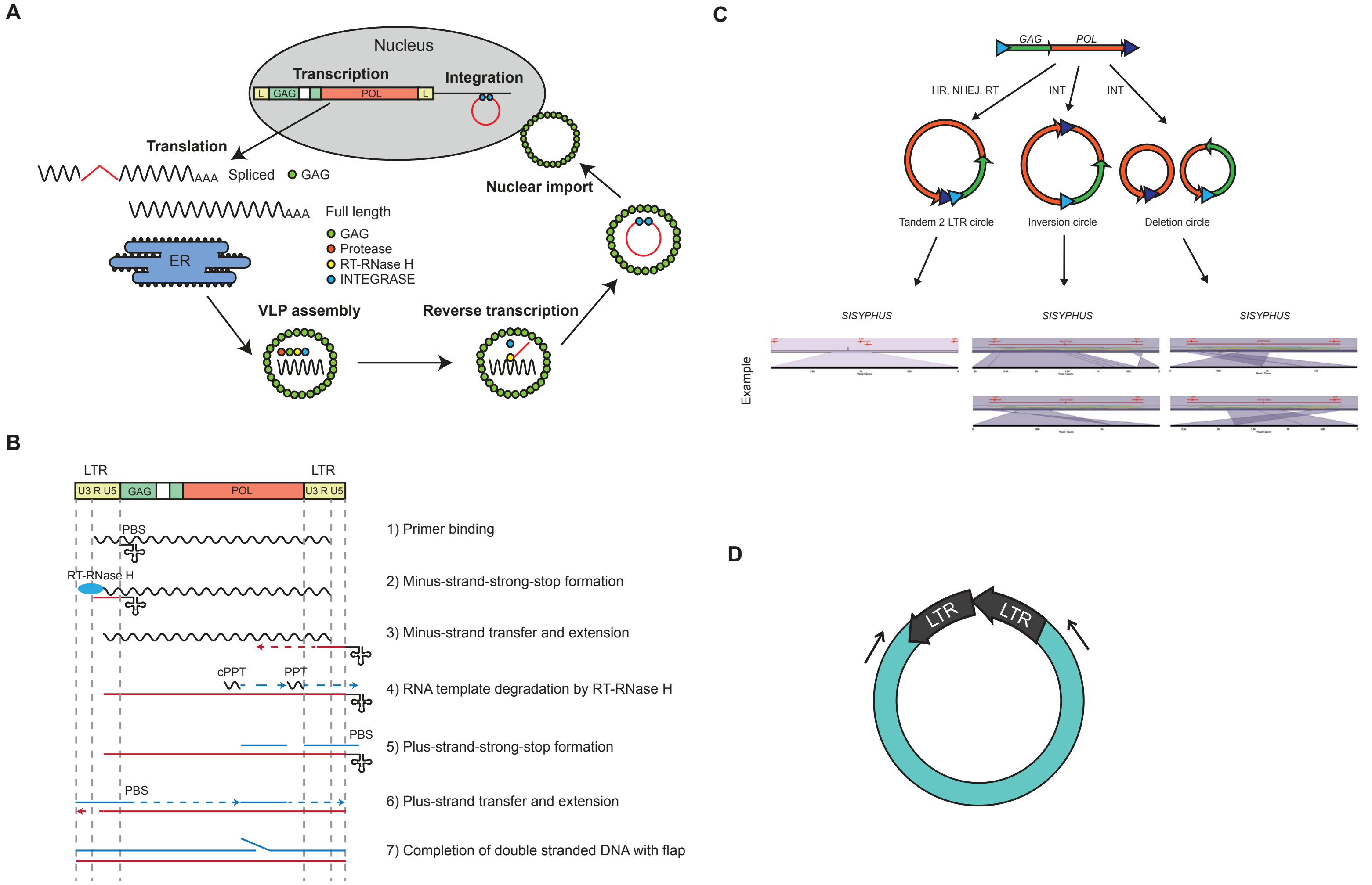


**Supplemental Figure S1. Replication cycle of LTR retrotransposon in plants.** (A) Production of virus-like-particles (VLPs) is initiated by transcription of functional elements. For *COPIA* elements in Arabidopsis, introns are spliced from subgenomic RNA to make only GAG proteins (Oberlin et al., 2017). Polyproteins encoded by full-length RNA comprise enzymes essential for reverse transcription and integration. After polyproteins are cleaved by protease, RT-RNase H processes full-length genomic RNA by reverse transcription. Double-stranded VLP DNA enters the nucleus and inserts into new genomic loci mediated by integrase. Abbreviations: L, LTR; ER, endoplasmic reticulum (B) Reverse transcription is initiated by tRNA primers that bind to the primer binding site (PBS). Minus-strand strong-stop DNA is made, transferred to the 3' LTR, and extended toward to 5' end. After removal of genomic RNA by RT-RNase H, RNA polypurine tract (PPT) fragments resistant to digestion are used for plus-strand strong-stop DNA formation. Plus-strand strong-stop DNA is transferred to 5' LTR and extended. Central PPTs (cPPT) produce additional plus-strand DNA that forms a flap structure by invasion of the extended plus-strand DNA from 5' end. (C) Examples of circular VLP DNA formation. Diagrams of circular VLP DNA formation (modified from Garfinkel et al., 2006) are shown to illustrate different types of products by homologous recombination (HR), nonhomologous end joining (NHEJ), reverse transcription (RT), or integrase (IN). Examples of Oxford Nanopore read alignments to *SISYPHUS* are shown for each circular DNA type using Ribbon (Nattestad et al., 2016). (D) Inverse PCR for circular DNA detection. Arrows indicate the orientations of two primers.
