## Supplemental Fig. S2 for "*Arabidopsis* LTR retrotransposons and their regulation by epigenetically activated small RNA"

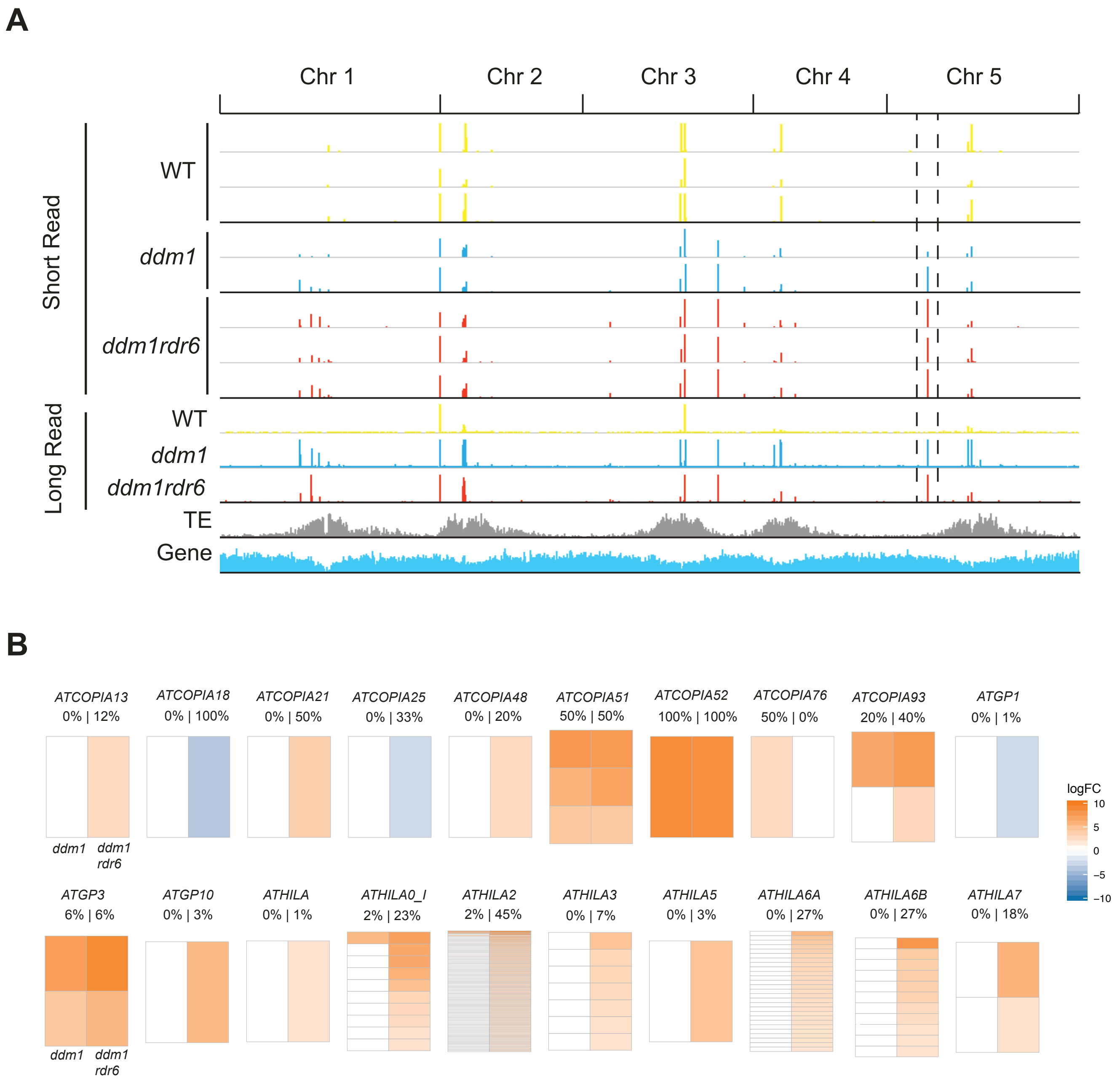


**Supplemental Figure S2. Genome-wide analyses of VLP DNA products in *ddm1* and *ddm1rdr6*.** (A) 3 replicate libraries of short sequencing reads (Illumina) and long sequencing reads (Oxford Nanopore) from VLP DNA samples were aligned to the Arabidopsis genome. Peaks of read alignments unique to *ddm1* and *ddm1rdr6* correspond to LTR retrotransposon loci that produce VLP DNA, whereas wild-type (WT) samples share DNase-insensitive background regions. Dashed lines indicate *EVADE*. Density of transposable elements (TE) and genes is shown. (B) Heat maps of log2 fold changes of VLP DNA enrichment in *ddm1* and *ddm1rdr6* for LTR retrotransposons. Each *ddm1* and *ddm1rdr6* genotype was compared to WT to calculate fold changes from short-read replicate samples. The percentages of significantly enriched TEs within the family for each genotype are shown above the heatmaps.
