## Supplemental Fig. S3 for "*Arabidopsis* LTR retrotransposons and their regulation by epigenetically activated small RNA"

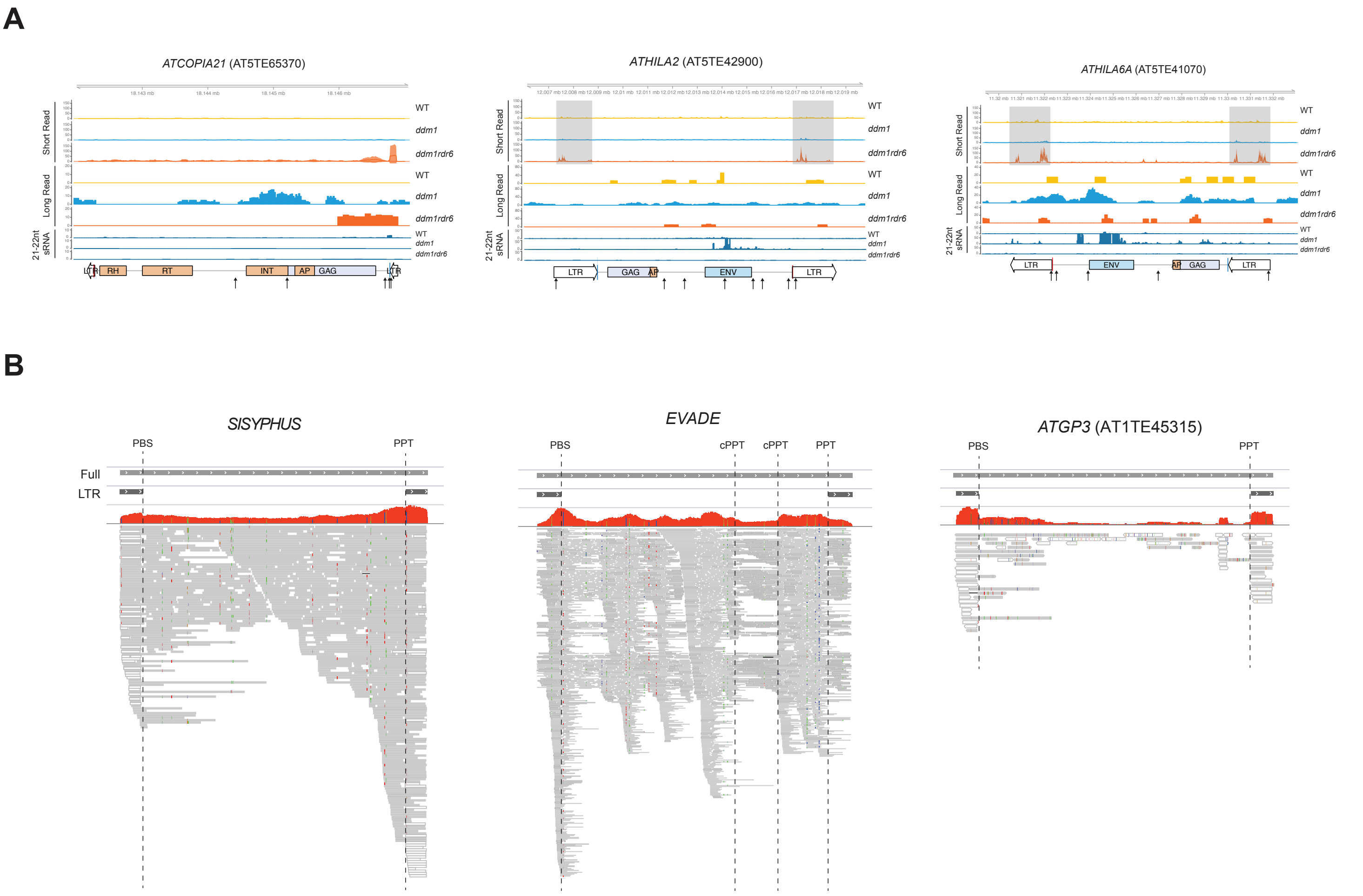


**Supplemental Figure S3. Additional VLP DNAseq data.** (A) Representative LTR retrotransposons significantly enriched in VLP DNA of *ddm1rdr6* plants. Short read (Illumina) VLP sequences were aligned to transposons, and mean read counts per million mapped and 95% confidence intervals of biological replicates are shown for wild-type (WT; yellow), *ddm1* (blue), and *ddm1rdr6* (orange) (n=3 for WT and *ddm1rdr6*; n=2 for *ddm1*). Gray shadow indicate the region showing partial VLP DNA from *ATHILA* elements. Long read sequence coverage was determined in a similar way from pooled VLP DNA replicates for each genotype and sequenced as one replicate by the long read sequencing platform. Annotation, small RNA, PBS and PPT sites are as in Figs. 1B, 6B. Target positions of miRNAs are indicated as arrows (see Supplemental Table S4 for details). (B) Alignments of long sequencing reads from VLP DNA in *ddm1rdr6*. Alignments are shown for *SISYPHUS* (AT3TE76225), *EVADE* (AT5TE20395), and *ATGP3* (AT1TE45315) VLP DNA. PBS, PPT, and cPPT positions are indicated as dashed lines relative to full and LTR annotation. Sequencing gaps and mismatches are indicated as in Fig. 3.
