## Supplemental Fig. S4 for "*Arabidopsis* LTR retrotransposons and their regulation by epigenetically activated small RNA"

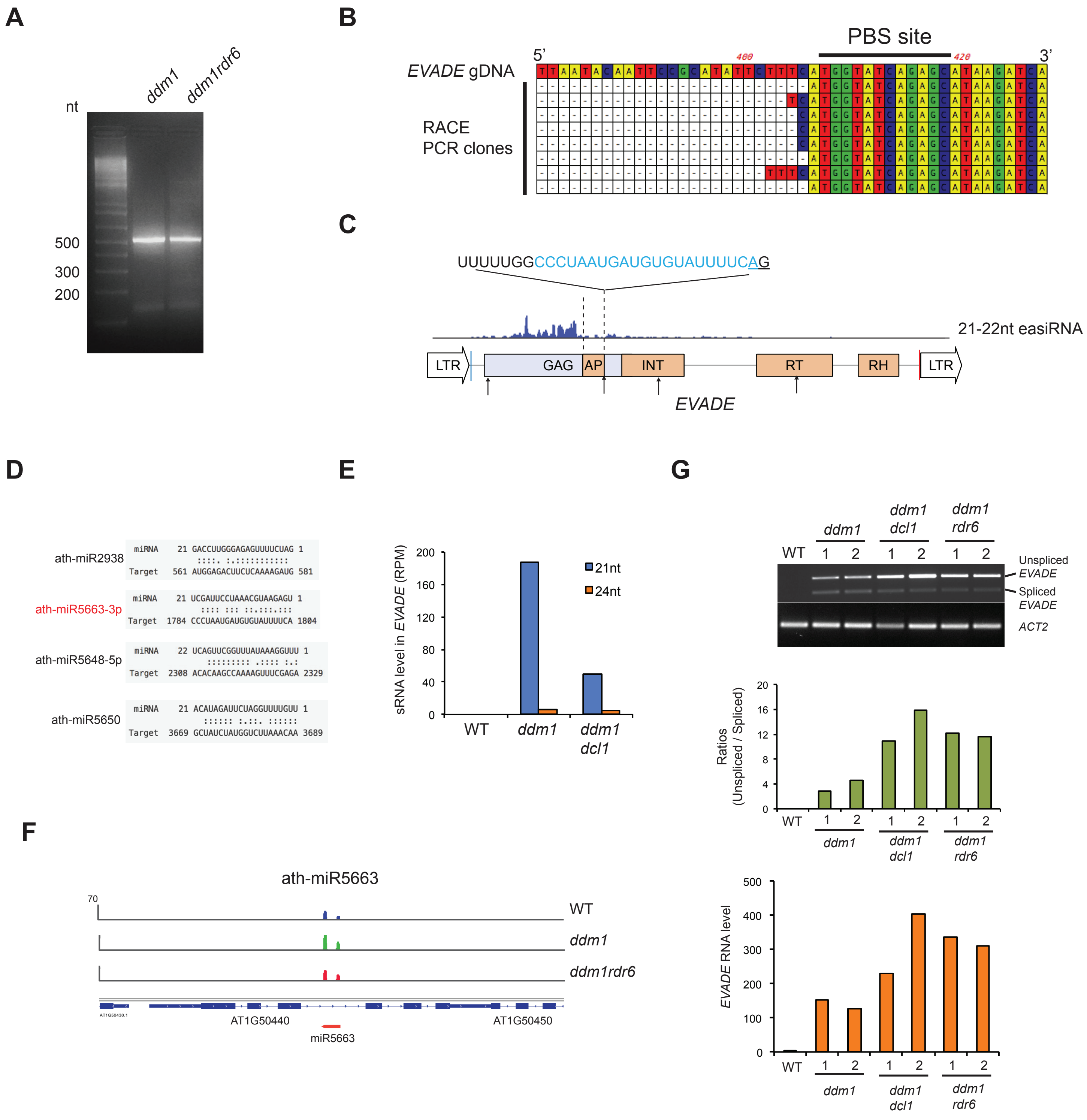


**Supplemental Figure S4. MicroRNAs targeting GAG and integrase (IN) regions of *EVADE*.** (A) 5' RACE PCR products from *ddm1* and *ddm1rdr6* using primers in the *EVADE* GAG gene. (B) 5' end sequences from the RACE products of *ddm1* (n=8) indicate RNaseH cleavage immediately upstream of the PBS. (C) miRNA target sites predicted by psRNATarget (Dai et al., 2018) are shown as arrows. 21-22nt easiRNAs and *EVADE* annotation are as in Fig. 1. Dashed lines indicate the intron of *EVADE*. The underlined and blue letters indicate the splicing acceptor AG nucleotide sequence and the target sequence of ath-miR5663-3p, respectively. (D) Sequence alignments of miRNAs with *EVADE* as a target sequence. (E) *EVADE* easiRNA abundance in reads per million (RPM) in wild-type (WT), *ddm1*, and *ddm1dcl1* mutants using a public dataset (GSE52951) (Creasey et al., 2014). (F) ath-miR5663 is detected in inflorescence tissues of wild-type (WT), *ddm1*, and *ddm1rdr6* (Creasey et al., 2014). (G) RT-PCR of *EVADE* for detection of spliced and unspliced forms in *ddm1*, *ddm1dcl1*, and *ddm1rdr6* (Top panel). Ratios of unspliced/spliced were calculated using band intensity from RT-PCR. *EVADE* RNA levels normalized by *ACT2* were obtained using RT-qPCR for each genotype (Bottom panel).
