## Supplemental Fig. S5 for "*Arabidopsis* LTR retrotransposons and their regulation by epigenetically activated small RNA"

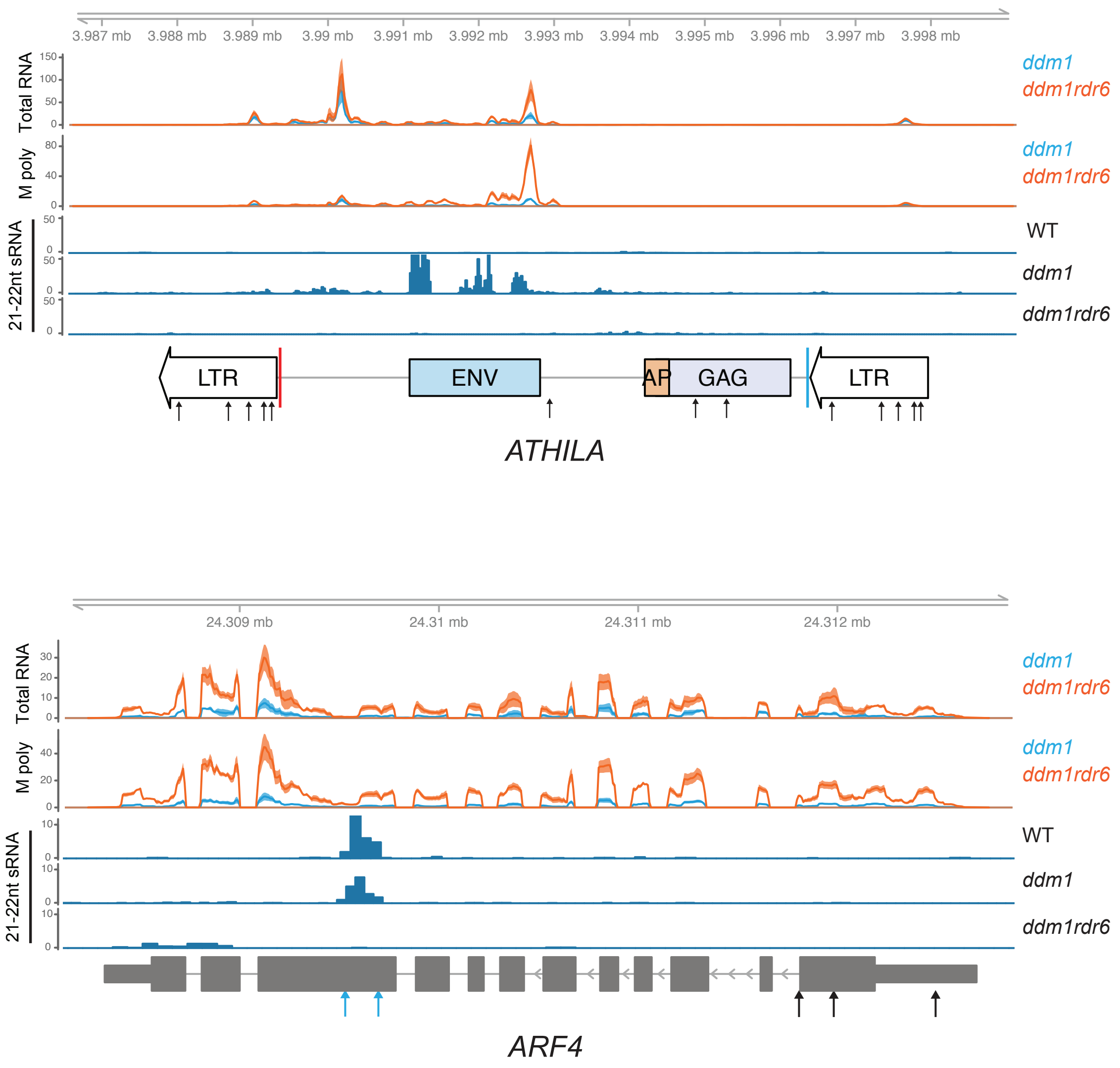


**Supplemental Figure S5. Polysomal occupancy of mRNA in *ddm1* and *ddm1rdr6*.** Total RNA and microsome-polysomal RNA (M poly) tracks are shown for *ATHILA* (AT4TE17360) and *ARF4* (AT5G60450), with annotation and 21-22nt easiRNA are as in Fig. 1. Black arrows indicate predicted miRNA target sites (Supplemental Table S4). Two target sites of the transacting small interfering RNA, tasiR-ARF4 are indicated as blue arrows (Williams et al., 2005).
