## Supplemental Fig. S6 for "*Arabidopsis* LTR retrotransposons and their regulation by epigenetically activated small RNA"

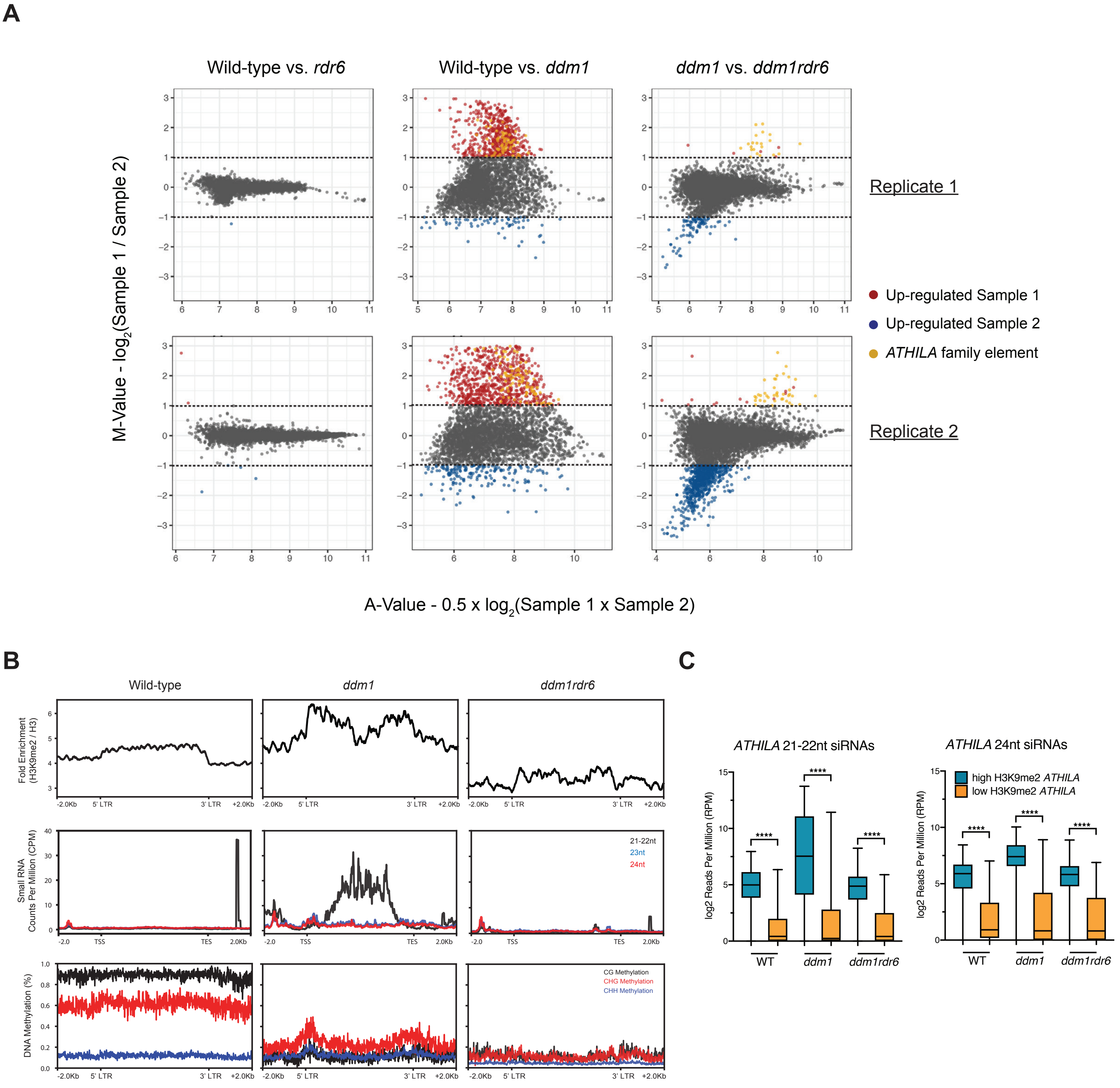


**Supplemental Figure S6. Meta-analysis of H3K9me2, small RNAs, and DNA methylation.** (A) MA plots of H3K9me2 ChIPseq data for wild-type, *ddm1*, and *ddm1rdr6*. Pairwise comparisons between genotypes revealing global changes in H3K9me2. log2 fold changes between Sample 1 and Sample 2 (M-value) are plotted against average read density between samples (A-value). Points represent ChIP-seq peaks that are up-regulated in Sample 1 (red), up-regulated in Sample 2 (blue), or not significantly different (gray). Annotated *ATHILA* family elements that are differentially regulated between samples are highlighted in yellow. (B) Metaplots depicting H3K9me2, small RNAs, and DNA methylation levels at the 73 most significantly affected *ATHILA* family elements across wild type, *ddm1*, and *ddm1rdr6* genotypes. Small RNA and DNA methylation data were obtained from a previously published study (Creasey et al., 2014). (C) Direct comparison of small RNA abundances between *ATHILA* family elements that gain RDR6-dependent H3K9me2 in *ddm1* and those that do not. Both 21-22nt and 24nt small RNA levels are shown in log2 reads per million (RPM) across wild-type (WT), *ddm1*, and *ddm1rdr6* genotypes. A two-sample Kolmogorov–Smirnov test was used to compare differences between *ATHILA* classes (**** p-value < 0.0001).
