## Supplemental Fig. S7 for "*Arabidopsis* LTR retrotransposons and their regulation by epigenetically activated small RNA"

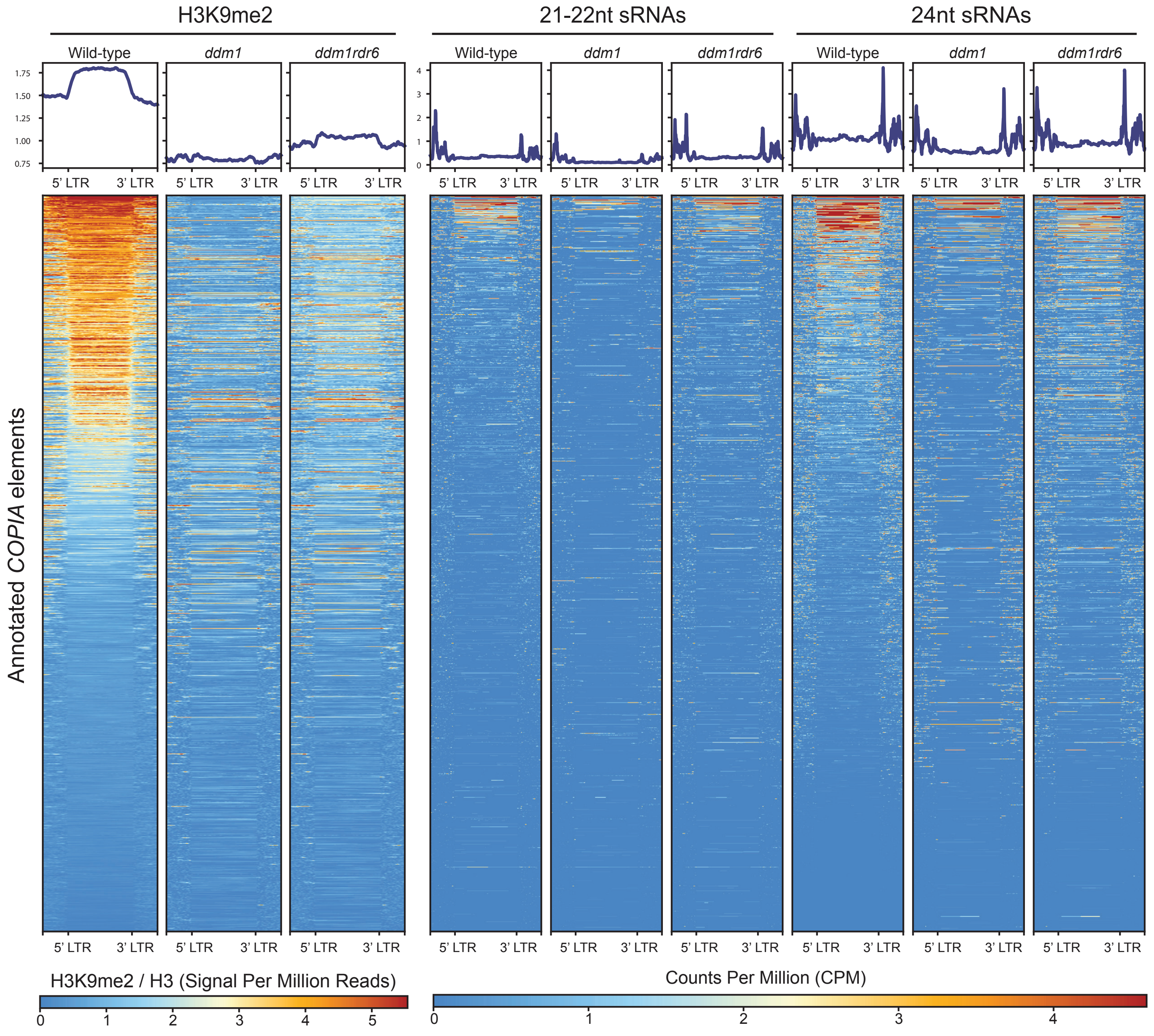


**Supplemental Figure S7. Up-regulation of H3K9me2 at *COPIA* family loci in *ddm1rdr6* as compared to *ddm1*.** H3K9me2 signal at transposable elements from multiple *COPIA* families was analyzed across wild-type (WT), *ddm1*, and *ddm1rdr6* genotypes and correlated with previously published small RNA data (Creasey et al., 2014).
